# Role of the metallo-reductase FADING and vacuolinos in anthocyanin degradation in flowers and fruits

**DOI:** 10.1101/2023.12.21.569356

**Authors:** Valentina Passeri, Shuangjiang Li, Pamela Strazzer, Enric Martínez i Calvó’, Esther Senden, Flavia Guzzo, Mauro Commisso, Cornelis Spelt, Michiel Vandenbussche, Mattijs Bliek, Walter Verweij, Ronald Koes, Francesca M. Quattrocchio

## Abstract

Anthocyanins are vacuolar pigments that confer red-violet colors to plant tissues. Pigmentation patterns result from spatio-temporally regulated anthocyanin synthesis and degradation. Mutational inactivation of a conserved MYB-bHLH-WDrepeat-WRKY transcriptional complex (MBWW) causes degradation of anthocyanins and ‘fading’ of flower color via a pathway that involves *FADING (FA)*. Here we show that *FA* encodes a vacuolar membrane Fe-reductase-oxidase that promotes anthocyanin degradation. In wild type petals anthocyanins in the central vacuole (CV) are stable, because FA-GFP is upheld in small vacuoles (vacuolinos) and kept away from the CV, indicating that vacuolinos act as gatekeepers in protein trafficking. In cells lacking vacuolinos, including *mbww-* mutant petals, FA-GFP reaches the CV and triggers anthocyanin degradation. Virus-induced gene silencing (VIGS) of an FA-homolog in pepper fruits prevented the “fading” of anthocyanins during fruit maturation. These findings provide new insights to breed ornamental and food crops with increased anthocyanin-content and enhanced nutritional value of edible parts.

## Introduction

The red and purple color of most flowers, leaves, and fruits results from the accumulation of anthocyanin pigments in the central vacuoles (CV) of (sub) cells. Pigmentation patterns are thought to be specified by the expression patterns of transcription factors that activate structural anthocyanin genes, encoding enzymes of the pathway in specific tissues^1^. Indeed, ectopic expression of these transcription factors is in many cases sufficient to induce the accumulation of anthocyanins in tissues where they are normally lacking^2–5^, which has health promoting effects in edible crops^6^.

Many fruits and flowers accumulate anthocyanins early in development that disappear in later stages, indicating that anthocyanins are not as stable as often assumed, and that their turnover also has a role in establishing a pigmentation pattern. While the synthesis of anthocyanins is one of the best studied biosynthetic pathways in plants, very little is known about their degradation and turnover. Petals of *Brunfelsia calycina* lose anthocyanins after bud opening and express a peroxidase that can trigger anthocyanin degradation in vitro^7^. In vivo evidence (e.g., from mutants) is currently lacking. In petunia, genetic factors required in vivo for the fading of petals are known for four decades^8^, but the underlying mechanisms have remained obscure^5^.

The coloration of petunia petals results from the accumulation of anthocyanins in the large central vacuole of epidermal cells, while underlying mesophyll lacks anthocyanins. The transcription of structural anthocyanin genes is driven by a complex (MBW) containing the MYB protein ANTHOCYANIN2 (AN2), the bHLH AN1 and the WD-repeat AN11. A related complex (MBWW) containing AN1, AN11, the MYB protein PH4 and the WRKY protein PH3 activate downstream genes, *PH1* and *PH5*, encoding vacuolar P-ATPase that acidify the vacuolar lumen conferring a reddish violet petal color. The same MBWW complex activates genes, including *PH1* and a small *Rab5a GTPase*, involved in the formation of vacuolinos, which are small (1-10 μm) vacuoles that coexist with the large central vacuole (CV) where anthocyanins are stored^9^ and act as intermediate stations for newly synthesized vacuolar proteins on their way to the CV^9,10^. Interestingly, this complex is required for stability of the anthocyanins in the central vacuole, as inactivation of MBWW triggers “fading” of the petal color.

The same MBWW is required for the acidification and the stability of anthocyanins in the vacuole^8,11–13^. In these mutants the anthocyanins fade after petal opening thereby shifting petal color from blue-violet to white^8,11^ (Fig. 1a; Supplementary Fig. 1a-b). Fading is suppressed by mutations blocking the decoration of anthocyanins with sugar and acyl groups and by a recessive allele of *FADING*^8^. Fading is nearly complete for highly substituted anthocyanin, and only partial, or even absent, in flowers accumulating more simple anthocyanins (ref^8^ and Supplementary Fig. 1a-b).

**Figure 1.**
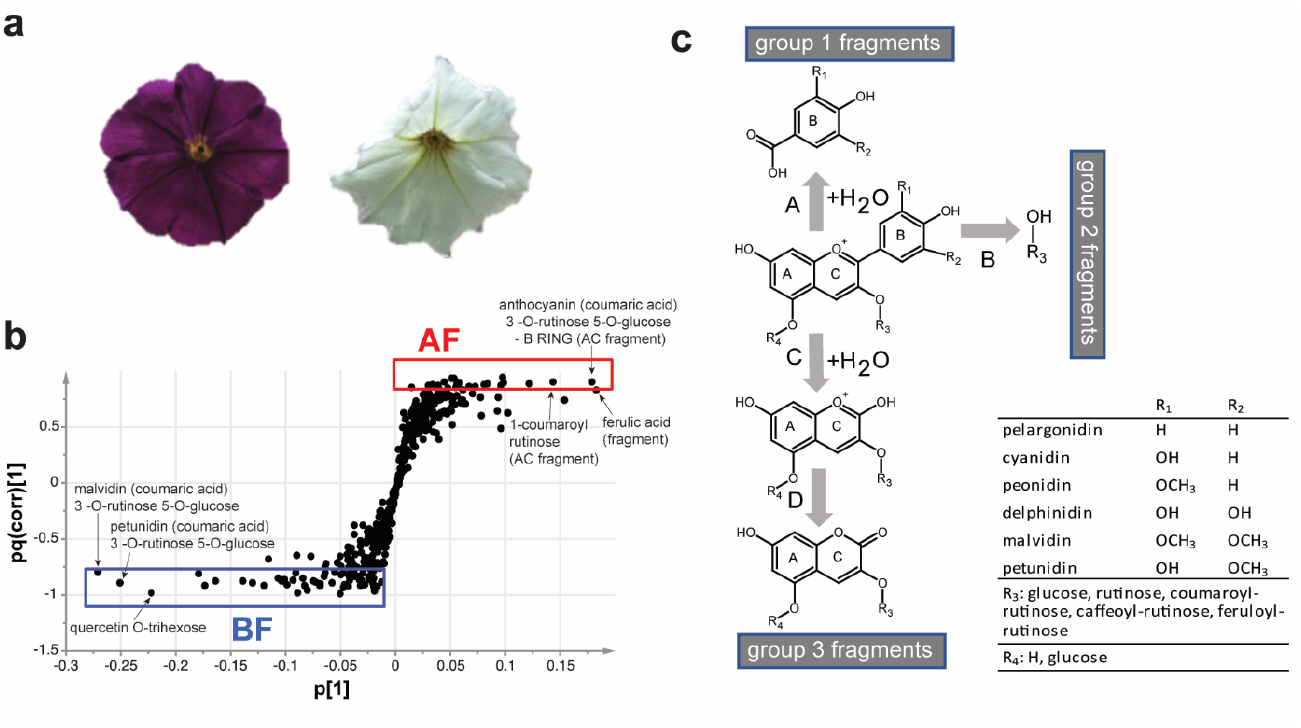
Identification of the *FADING* gene. **(a)** Flower of the *Petunia hybrida* F1 hybrid V74XR149 just after bud opening (left) and 6 days later when pigment fading is completed. **(b)** S-loading plot of V74xR149 flowers before (BF) and after fading (AF), obtained by mapping p against pq(corr) in OPLS-DA. Each point represents a metabolite detected and quantified by LC-MS. p is an indicator of the “weight” of each molecule in the model, which directly depends on its abundance; pq(corr) indicates the ability of a metabolite to distinguish the samples before and after fading (two classes). Anthocyanins are shown to accumulate specifically before fading, while putative anthocyanin fragments characterized flowers after fading. The complete dataset is reported in DataSheet 1. **(c)** Putative fragmentation of anthocyanin molecule during fading.

To study the mechanism of fading in petals, we identified the petunia *FA* gene by transposon tagging and found that it encodes a ferric-reductase-oxidase (FRO) expressed in tissues that synthesize anthocyanins (e.g., petal epidermis) and acyanic tissues (e.g., petal mesophyll and leaves). In wild type epidermal petal cells, FA-GFP is upheld in vacuolinos and released from the membrane into the lumen, not reaching the CV. Instead, in mutants that lack vacuolinos (*ph3* and *ph4*) FA reaches the CV where it triggers the degradation of anthocyanins.

We report here a first application of these findings for the breeding of bell pepper cultivars with persistent anthocyanin pigmentation.

## Results

### Characterization of the fading process

Petals of petunia *ph4* mutants have a bluish (‘blue violet’) colour at bud opening, as vacuolar acidification is impaired, and subsequently ‘fade’ to white in 4-6 days (Fig. 1a). To characterize fading in more detail, we analysed petal tissue before fading (stage 7) and after (stage 8) by HPLC-ESI-IT, followed by multivariate analysis (OPLS-DA). This separates stage 7 and stage 8 flowers based on their metabolite content. The S-loading plot [p *versus* pq(corr)] showed that anthocyanins (i.e., glycoside derivatives of malvidin and petunidin acylated with coumaric acid) are characteristic for petals before fading, while other compounds are specific for petals after fading (Fig. 1b). Some of the latter compounds are putative anthocyanin degradation products, based on their m/z value and fragmentation pattern, and validated by high-resolution UPLC-ESI-QTOF analysis (Supplementary Table 1). The putative anthocyanin fragments can be classified into three different groups (Fig. 1c). Fragments in group 1 exhibit carboxylated and glycosylated B rings. The formation of carboxylic acid of B ring following C ring fragmentation was previously described for anthocyanins^14^ and other flavonoids^15^. Group 2 fragments contain glycosylated hydroxycinnamic acid, and most likely originate from the 3-O-chain of the acylated anthocyanin, while group 3 fragments, which contain the decorations R3 and R4 on A and C rings, most likely result from disruption of the bond between C-ring and B-ring, accompanied by water addition.

This fragment is mainly detected in the negative ionization mode in MS, supporting a rearrangement of the C ring and loss of the positive charge typical of the flavylium ion. Similar anthocyanin byproducts have been described for anthocyanidin-3,5-diglucosides undergoing structural transformations which lead to the unexpected formation of A+C ring-derived glucosides^16^. Several other secondary metabolites were specifically found before (e.g., quercetin O-trihexose) and after (ferulic acid fragment) fading (Fig. 1b), suggesting that other compounds beyond anthocyanins may undergo degradation during this process. The complete dataset is provided in DataSheet 1.

### Isolation of *FA* by transposon-tagging

Line R149 is an *FA*^+^ *ph4* line^12^ that contains like its parent, W138, ∼400 highly mobile *dTPH1*-type transposons^14^ and contains mutations that block hydroxylation (*hf1, hf2*) and rhamnosylation (*rt*). Hence, R149 petals accumulate cyanidin 3-glucosides, which are (relatively) insensitive to fading^8^. We crossed R149 with V74, a *ph4 fa HF1 RT* line that synthesizes fully substituted anthocyanins (malvidins) that do not fade (due to *fa)*. Among ∼7000 *FA*^+/v74^ *ph4ph4* progeny having blue-violet petals that faded, we found two mutants (plants R2033-1 and R2057-1) with blue-violet petals that did not fade, indicating that the R159-derived *FA*^*+*^ allele was inactivated (Fig. 2a). R2057-1 petals had occasionally spots that ‘faded’ to white in 3-6 days after bud opening, suggesting that the new *fa*^*R2057*^ allele is an unstable transposon-insertion allele. In R2033-1 petals, by contrast, we never observed such somatic reversions (Fig. 2a).

**Fig 2.**
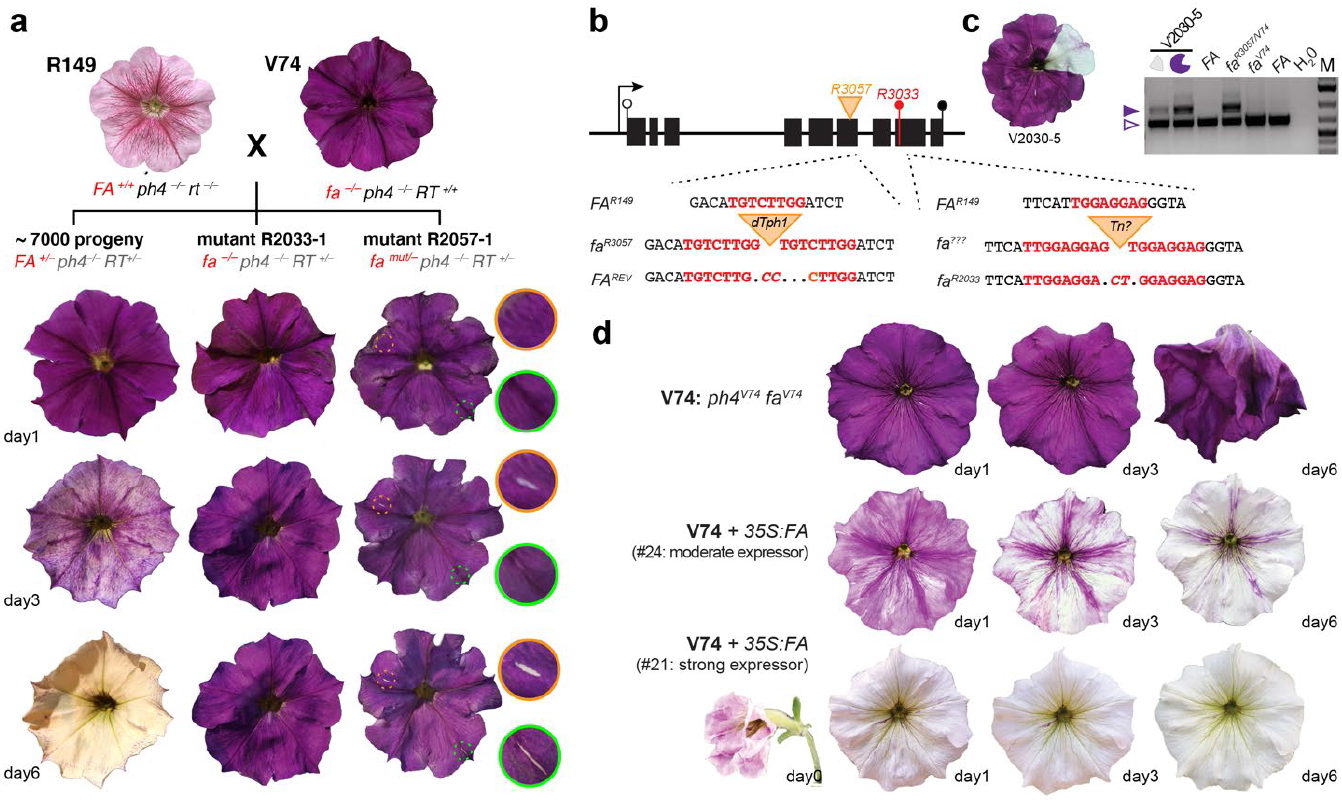
Identification of the *FADING* gene. **(a)** Isolation of mutant alleles *fa*^*R2057*^ and *fa*^*R2033*^ by targeted transposon mutagenesis. **(b)** Diagram of *FA* showing the position of introns and exons (black rectangles), start and stop codons (open and closed black lollipops respectively), transposon insertions (triangles) or footprints (red lollipop) in *FA* and mutant alleles. The lower parts shows sequence alterations in *fa*^*2033*^ and *fa*^*2057*^, derived somatic revertants (*FA*^*REV*^) and inferred transposon insertion progenitor alleles. Red font indicates target site duplications. Nucleotides lost or duplicated from the bottom strand after transposon excision are indicated by dots and italics respectively. **(c)** PCR analysis of the *fa*^*2057*^ region containing the *dTPH1* insertion in the violet *fa*^*R2057/v74*^ and white FA^REV/v74^ part of a flower individual (V2030-5) of (back)cross R2057-1 x V74. The closed and open violet arrowheads indicates PCR fragments with or without a *dTPH1* insertion **(d)** Phenotype of V74 flowers (*ph4 fa*) and transgenic sibling expressing *35S:FA* at moderate or high level.

Because “spontaneous” mutations in the W138 background are mostly due to *dTPH1* insertions ^12,13,17–20^, we sequenced flanking regions of (nearly) all *dTPH1* elements in R2033-1, R2057-1 and nine *FA*^+^ siblings of each. We identified a candidate *FA* gene that was in R2057-1 disrupted by a *dTPH1* insertion and in R2033-1 by an 8-bp insertion that probably arose by the insertion and excision of a transposon, whereas all their siblings lacked these mutations (Fig. 2b). We found no germinal revertants among ∼7000 V74xR2057-1 backcross progeny, but one plant (V2030-5) produced a flower with a large somatic *FA* revertant sector that could be analyzed by PCR (Fig. 2c). In the violet (*fa*) tissue the *dTPH1* insertion in *FA* candidate was maintained while it was severely reduced in the white tissue (*FA*^*rev*^) –but not fully abolished, as *fa*^*2057*^ is preserved in the underlying colorless mesophyll – and abundant excision product with a 6-bp footprint appeared (Fig. 2b-c).

Next, we generated transgenic V74 plants (*fa, ph4*) and found that expression of *FA* from the constitutive viral 35S promoter restored in the fading phenotype (Fig. 2d). In some transformants the petals were colored at bud opening and faded thereafter, while in flowers of transformants with higher transgene expression, remained nearly white at bud opening (Fig. 2d; Supplementary Fig. 1c). Expression of a GFP-FA fusion protein in V74 plants also restored the fading process (Supplementary Fig. 1d). Together, these results prove that the identified gene is encoded by *FA*.

### *FA* encodes a ferric-reductase-oxidase

FA belongs to the family of ferric-reductase-oxidase (FROs) metalloreductases, which are involved in iron and copper homeostasis^21–23^ and share similarity with yeast ferric reductase ScFRE1^24^, and HsGP91phox, a subunit of the human NADPH oxidase that produces reactive oxygen species (ROS) in response to pathogens^25^.

FRO proteins cluster in 4 distinct clades, each containing at least one protein from Arabidopsis and petunia (Fig. 3a). FA from *P. hybrida*, the *P. axillaris* N ortholog PanFA, and two closely related paralogs (PanFADING-LIKE1, PanFAL1 and PAnFAL2) cluster with FRO6 and FRO7 from *A. thaliana* in clade B, while the other Petunia and Arabidopsis paralogs cluster in the distinct clades. FA homologs are present in a broad range of plant species including *Solanaceae* and other dicots, the early dicot *Nelumbo nucifera*, monocots (e.g., *Oryza sativa* and *Zea mays*), basal angiosperms (*Amborella trichopoda, Cinnamomum micranthum*), gymnosperms (*Pseudotsuga menziesi*), lycophytes (*Selaginella moellendorffii*), mosses (*Physcomitrella patens*), liverworts (*Marchantia polymorpha*) (Fig. 3a). This indicates that the B (FA) clade of FRO proteins is of ancient origin.

**Figure 3.**
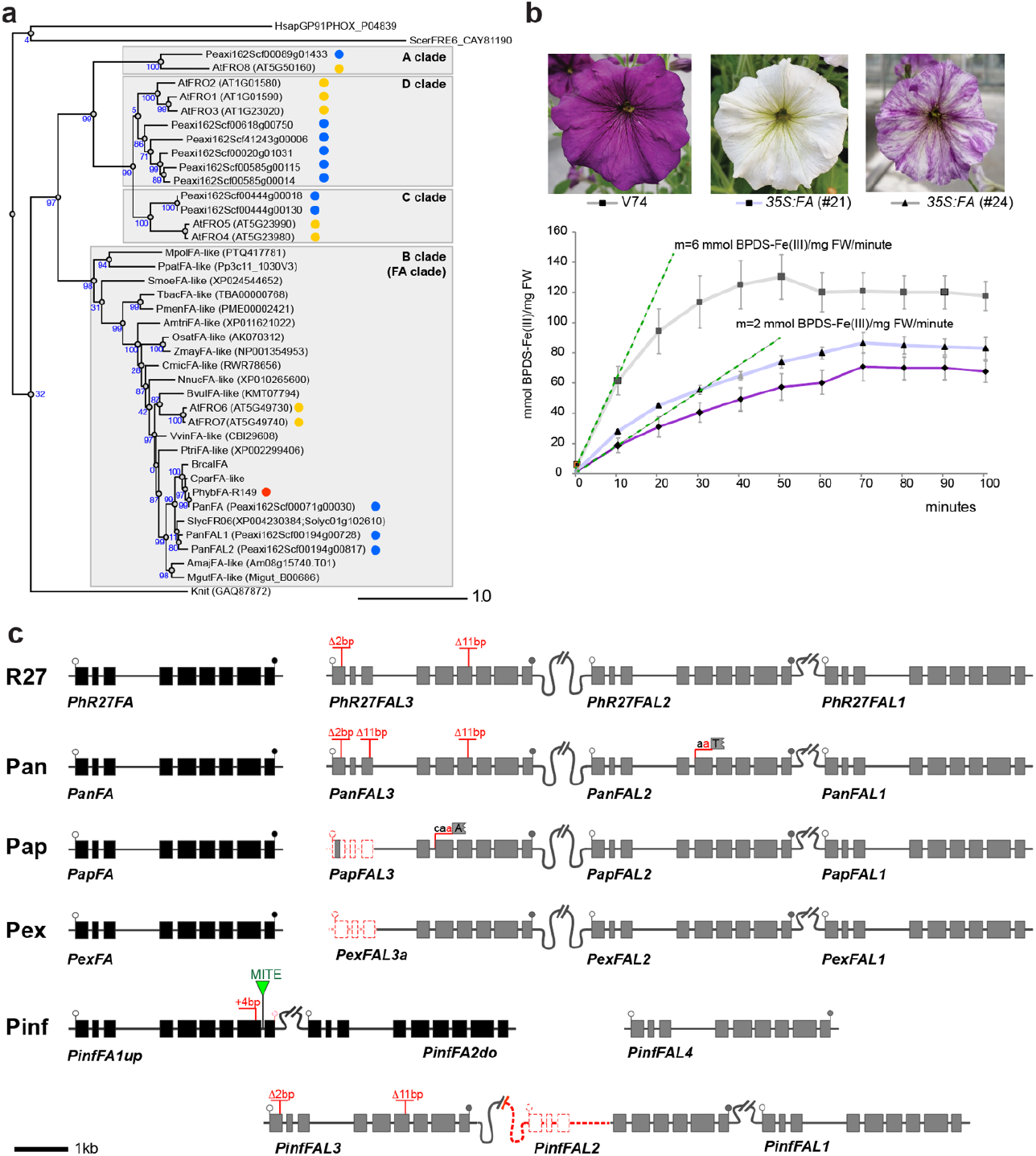
*FADING* encodes a ferric reductase oxidase of the FRO family. **(a)** Phylogenetic tree of FRO proteins from *Amborella trichopoda* (Amtri), *Arabidopsis thaliana* (At), *Antirrhinum majus* (Amaj), *Beta vulgaris* (Bvul), *Cinnamomum micranthum* (Cmic), *Calibrachoa parviflora* (Cpar), *Homo sapiens* (Hsap), *Oryza sativa ssp. Japonica* (Osat), *Klebsormidium nitens* (Knit), *Petunia axillaris* N (Pan or Peaxi), *Petunia inflata* (Pinf), *Physcomitrella patens* (Ppat), *Populus trichocarpa* (Ptri), *Pseudotsuga menziesii* (Pmen), *Marchantia polymorpha* (Mpol), *Mimulus guttatus* (Mgut), *Nelumbo nucifera* (Nnuc), *Saccharomyces cerevisiae* (Scer), *Selaginella moellendorffii* (Smoe), *Solanum lycopersicum* (Slyc), *Taxus baccata* (Tbac), *Vitis vinifera* (Vvin),and *Zea mays* (Zmay). Gene identification or Genbank accsion numbers are in paranthesis. Blue numbers on 1) How did you get a initial rate (tangent) for V74 = 1 mmol BPDS/mg FW according to this Fig. but seems to me closer to 2. 2) Sure that it is mmol, not micromol? branche (nodes) indicate SH-like aLRT branch support (x 100). **(b)** Fe^2+^ oxidation activity in petal extracts from V74 (*fa*) and two transgenic plants with moderate (#24) and high(#2) transgen expression and a fading phenotype (average ± SE, n=3).. The initial reaction rate is calculated from the tangent at t = 0. **(c)** Diagrams depicting the structure of *FA* and *FAL* homologs in *P. hybrida* R27, *P. axillaris* N (Pan), *P*.*axilaris* P, (Pap). *P. exserta* (Pex) and *P. inflata* (Pinf).

To assess whether FA has Fe oxidoreductase activity, as expected for a member of the FRO family^26,27^, we compared ferric-reductase activity in *fa* mutant petals (*fa ph4* line V74) and two transgenic lines for the *35S:FA* construct (V74 *35S:FA*) using the colorimetric Fe^2+^ indicator bathophenanthroline disulfonic acid (BPDS)^25^. *fa* petals contain low Fe-reducing activity, which was not unexpected as the petunia genome encodes multiple FRO proteins and possibly other Fe reducing activities. However, activity was >3-fold higher in isogenic *fa 35S:FA* petals showing full complementation (line #21) and much lower in petals with partial complementation (line #24) supporting that FA has Fe-reducing activity (Fig. 3b).

The genomes *P. hybrida* R27, an isogenic progenitor of R149, and progenitor species from which *P. hybrida* derived – *P. axillaris, P. exserta, P. inflata* – contain besides *FA* a cluster of three closely related genes, here named *FA-LIKE1* (*FAL1*), *FAL2* and *FAL3* (Fig 3c). *FAL3* acquired various inactivating mutations, making it a pseudogene across all accessions. FAL2 appears to follow suit, as the *P. inflata* allele acquired a substantial deletion, and *P. axillaris* N allele a potentially crippling splice site mutation. The *P. inflata* contains two FA orthologs (PinfFA1 and PinfFA2) that are ∼ 9kb apart and apparently result from a recent duplication of the gene body plus >7 kb flanking sequences. *PinfFA1* is likely to have become a pseudogene, despite its young age, due to a frameshift mutation (Fig. 3c).

### *FA* expression

In the *P. hybrida* M1xV30 and R27, the *FA*^*+*^ allele is expressed in petals reaching highest levels after bud opening (stage 5) when color fading sets in (Fig. 4a-c). Mutations in *AN1, PH3* or *PH4* have little or no effect on *FA* expression in the petal limb (Fig. 4b), excluding that MBWW stabilizes anthocyanins by inhibiting *FA* expression. *FA* is also expressed in tissues that lack anthocyanins, such as sepals, stems, mature leaves, and ovaries. Expression of the *FAL* genes in stage 4 petals of R27 is very low (PhR27FAL1) to undetectable (PhR27FAL2 and PhR27FAL3) compared to FA, explaining why mutation of *FA* is sufficient to block fading (Fig. 4c).

**Figure 4.**
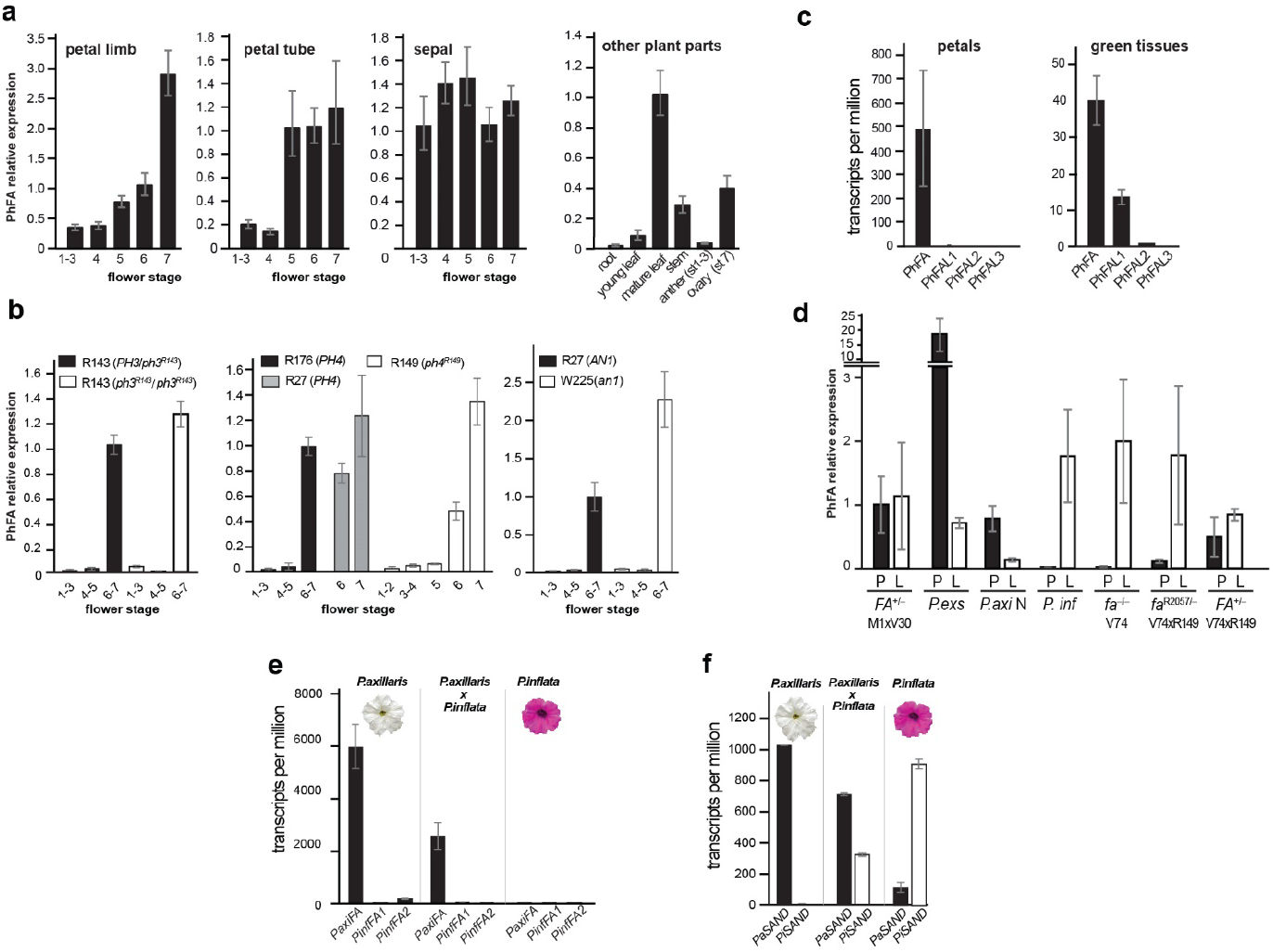
Expression of *FA* and *FAL* genes in *P. hybrida* R27 and ancestral Petunia species. **(a, b)** FA expression in organs from flowers of different stages, and other tissues of M1xV30 (a) and petals of line R27 an isogenic *an1, ph3* and *ph4* mutants lines **(b)** determined by qRT-PCR and normalized to RAN (mean ±SD, n = 2 biological replica and 4 (a) or 2 (b) technical replicates). **(c)** Expression of FA and FAL genes in petals and green tissues (leaves and/or shoots) of line R27, inferred from RNA seq data (mean± SD, n=2). **(d)** *FA* expression in leaves and petals of different P. hybrida lines and wild ancestors, determined by qRT-PCR and normalized to RAN (mean ±SD, n = 3 biological replica and 2 technical replicates). **(e**,**f)** Expression of *FA* and *SAND* alleles in petals of *P. axillaris* N, *P. inflata* and F1 progeny inferred from RNAseq data. Expression is given as number of reads corrected for the length of the transcript.

The gene body of the recessive *fa*^*V74*^ allele does not contain obvious disruptive mutations and the encoded protein is highly similar to FA^R27^ (Supplementary Fig. 2) and is in leaves expressed at a similar level as *FA*^*+*^. However, *fa*^*V74*^ is in petals expressed at a very low level, explaining why this allele does not trigger flower color fading (Fig. 4d). To reveal the origin of *FA*^*+*^ and *fa*, we analyzed expression of *FA* alleles in the parental species *P. axillaris*, a moth-pollinated species with white petals, *P. exserta* a bird-pollinated species with red petals^28^, and *P inflata*, a bee-pollinated species with violet petals.

To assess whether the divergent FA expression in maturing *P. axillaris* and *P. inflata* petals results from cis- or trans-regulatory differences, we analyzed public RNA-seq data ^28,29^ from petals of *P. axillaris, P. inflata* and a F1 progeny and mapped reads for *FA* to the corresponding *P. axillaris* and *P. inflata* alleles and, as a control, *SAND*^30^. *SAND* mRNA is expressed at similar levels in petals of both species and the F1 expresses the *P. axillaris* and *P. inflata* alleles at similar levels (Fig. 4e-f). The *FA* reads from the petals of F1 plants, by contrast, map almost entirely to the *P. axillaris* allele, and there is little or no reads from the *P. inflata* allele (Supplementary Fig. 3). Therefore the differential expression of *FA* in *P. axillaris* and *P. inflata* petals is caused by a cis-regulatory difference, and (ii) the *FA* and *fa* alleles in *P. hybrida* most likely reflect introgressions of wild progenitor species.

#### Intracellular localization and trafficking of FA-GFP

In Arabidopsis, distinct FRO proteins reside in different subcellular compartments^21^, AtFRO5 and AtFRO2 reside in the plasma membrane for uptake of extracellular ferric iron (Fe^3+^), and possibly copper (Cu^2+^) by roots^21,31^, whereas AtFRO3 and AtFRO8 reside in mitochondria^32^. The FA ortholog *At*FRO7 localized in chloroplasts to facilitates iron import for chlorophyll assemblage, while AtFRO6 was reported to be on the plasma membrane^27^.

We transiently transformed petals by agroinfection to express FA proteins with an N- or C-terminal GFP-tag encoded by chimeric genes containing the FA coding sequence from the gene (gFA-GFP and GFP-gFA) or cDNA (cFA-GFP and GFP-cFA) driven by the viral 35S promoter. Immunoblot analysis showed that these petals expressed full-size fusion protein and little or no cleavage products (Supplementary Fig. 4a). As proteins expressed from *35S:cFA-GFP* were the most abundant and could in transgenic plants restore fading in *fa* flowers (Supplementary Fig. 1c), we used this construct, hereafter named *35S:FA-GFP*, for further experiments.

In petal mesophyll protoplasts or in leaf cells transiently expressed vacuolar proteins (e.g., PH5-GFP) reach the CV within 24 hours^33^. However, in protoplasts from the petal epidermis – distinguishable by the anthocyanins in their CV – vacuolar proteins arrive after 24 on vacuolinos, and need another 20-24 hours to reach the CV^9^.

In wild type petal mesophyll protoplasts, FA-GFP arrives on the tonoplast of the CV within 24 hours where it remains after 48 hours (Fig. 5a). In petal epidermis protoplasts, FA-GFP arrived after 24 hours in the membrane of vacuolinos and, surprisingly, in their lumen (Fig. 5a). After 48 hours, fluorescence in the lumen had increased, while little of no FA-GFP had moved on to the CV (Fig. 5a). Similar localization of FA-GFP was observed in petals of transgenic plants (Supplementary Fig. 4b). Thus, in petal epidermis, trafficking of FA-GFP is remarkably different from that of other tonoplast proteins, as FA-GFP is upheld in vacuolinos and relocated from the membrane to the lumen, whereas other vacuolar proteins, like PH5-GFP, move on to the CV.

**Figure 5.**
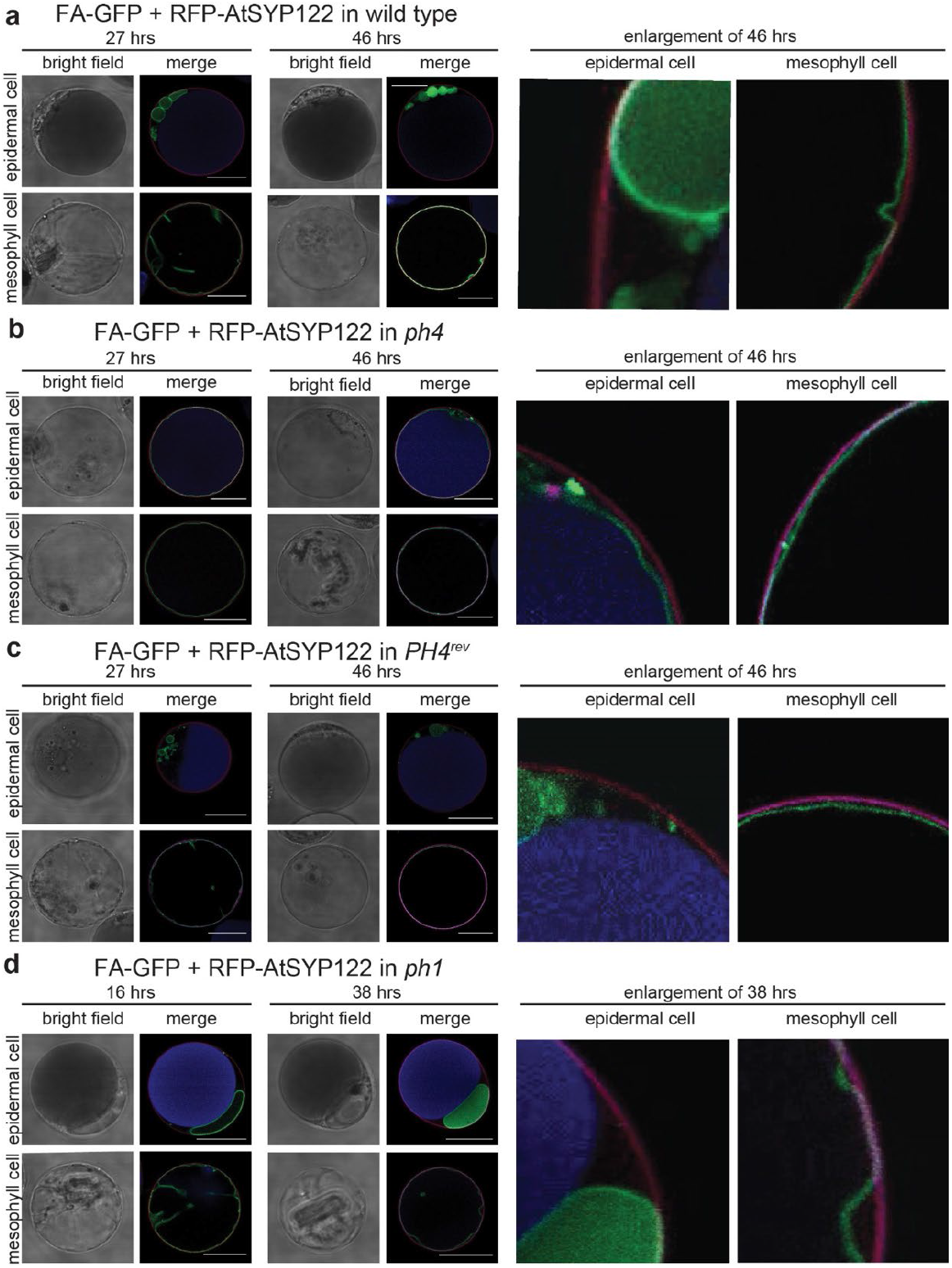
Subcellular localization of FA in transiently transformed petal protoplasts. **(a-d)** Localization of FA-GFP in protoplasts from petal epidermis and mesophyll of wild type **(a)**, *ph4* mutant **(b)**, *PH4* revertant **(c)** and *ph1* mutant **(d)** one or two days after transformation. RFP-AtSYP122 (visible in red) is used as marker of the plasma membrane, anthocyanins in the lumen of the CV of epidermal cells are visible in blue. The size bars equal 20 μm.

Mutation of *PH3* or *PH4*, which abolishes vacuolinos ^9,10^, affected trafficking of FA-GFP in epidermal cells, now moving directly (within one day) to the tonoplast of the CV^9,10^ (Fig. 5b). In mesophyll protoplasts of these mutant petals, FA-GFP behaves as in wild type and this sorting was the same in epidermal cells (Fig. 5b and Supplementary Fig. 4c). In epidermal petal protoplasts of a germinal PH4 revertant (*PH4*^*Rev*^) FA-GFP localized to the vacuolino membrane 24 hours after transformation, while after 48 hours GFP fluorescence arrived in their lumen but not on the CV (Fig. 5c), like wild type (Fig. 5a).

In epidermal petal cells of *ph1* mutants, protein trafficking from vacuolinos to the CV is strongly reduced and vacuolinos are enlarged^9^. In these *ph1* cells FA-GFP is visible on the membrane of the large vacuolinos one day after transformation and from where it gradually disappears into the lumen (Fig. 5d). In the mesophyll cells of the same *ph1* petals, FA-GFP localizes on the CV tonoplast already 16 hours after transformation where it remains later on (Fig. 5d).

To examine whether FA-GFP and PH5-GFP label different populations of vacuolinos, we co-expressed them in wild type petunia petal protoplasts. In 24 hours, FA-RFP and PH5-GFP co-localized to the same population of vacuolinos (Fig. 6a). PH5-GFP fluorescence remained on the membrane, suggesting that the internalization is specific for the FA-fusion proteins. When these constructs were co-expressed in epidermal petal cells lacking vacuolinos (e.g., *ph4* mutant line V74), both FA-RFP and PH5-GFP reached the tonoplast of the CV within 24 hours (Fig. 6b).

**Figure 6.**
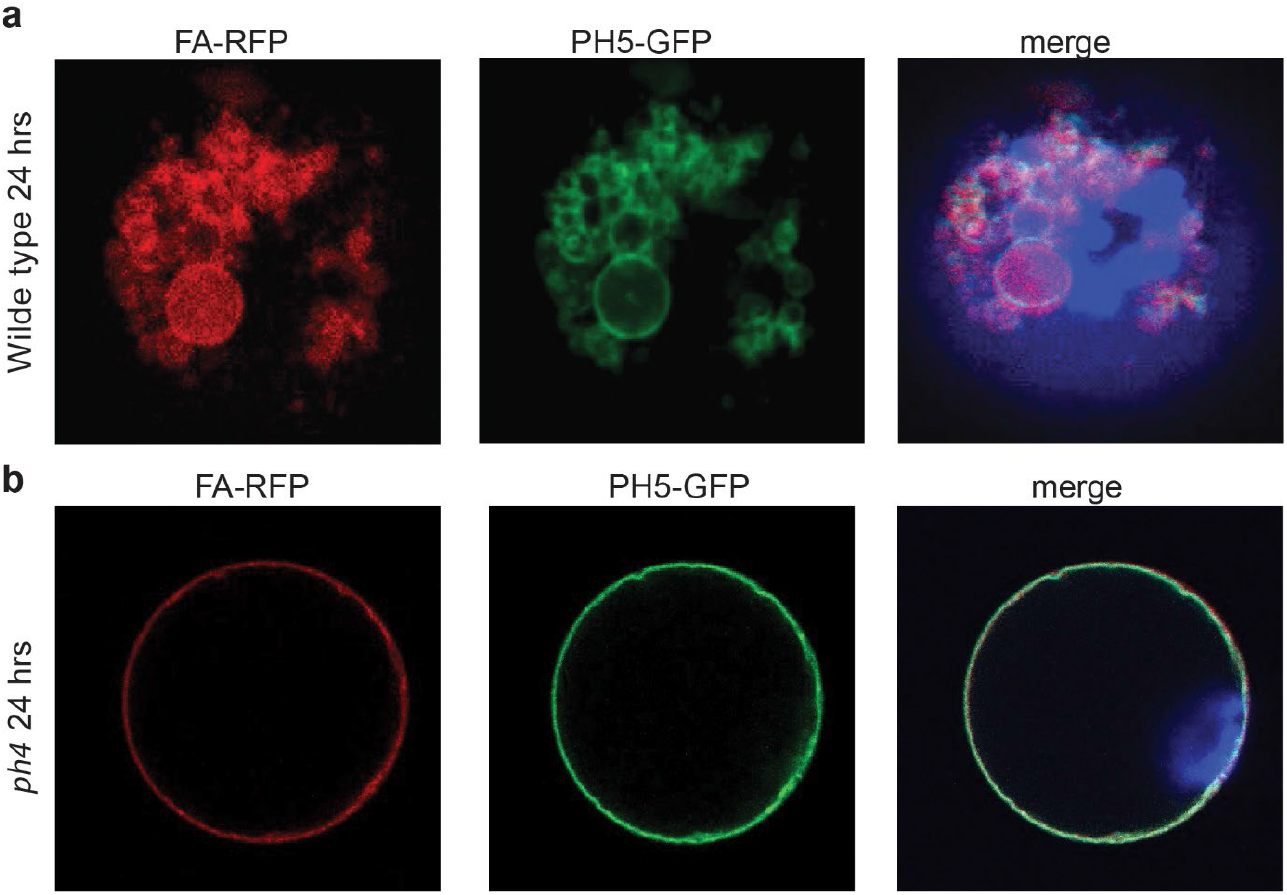
FA-RFP and PH5-GFP localize on the same population of vacuolinos. **(a)** In protoplasts from petal epidermis of the wild type M1×V30 F1 hybrid, one day after transformation, FA-RFP is visible as red fluorescence in the lumen of the vacuolinos which is marked on their membrane by PH5-GFP. **(b)** In protoplasts from petal epidermis of the *ph4* mutant line V74, both FA-RFP and PH5-GFP reach the tonoplast of the CV already one day after transformation.

### Localization of FA homologs and paralogs

To assess FAL proteins may be functionally similar to FA, we studied the intracellular localization of FAL1, the only clearly functional active of the FAL paralogs. We found that the trafficking of PanFAL1 in petal protoplast is indistinguishable from that of FA (Supplementary Fig. 5).

The vacuolar localization of FA and FAL1 is remarkable as the Arabidopsis homologs AtFRO6 and AtFRO7 had been localized in the plasma membrane and chloroplasts respectively^25^. To uncover the (evolutionary) origin of this difference, we swapped GFP-fusions of these proteins between Arabidopsis and petunia. In leaf cells of Arabidopsis and petunia, both AtFRO6-GFP and AtFRO7-GFP are observed on the tonoplast 24 hours after transfection (Supplementary Fig. 5a-b and 6a-b). Additionally, AtFRO7-GFP shows partial localization to chloroplasts (Supplementary Fig. 6a-b). In petunia protoplasts of the petal epidermis, both AtFRO6-GFP and AtFRO7-GFP localize to the vacuolino lumen, similar to FA-GFP, while in petal mesophyll cells, they reach the tonoplast within 24 hours (Supplementary Fig. 6 and Fig. 7).

These subcellular localizations suggest that members of the FRO family that cluster in the FA clade, all are vacuolar ferric-reductase-oxidases.

### Fading in other Solanaceae

*Brunfelsia* is a nightshade that is closely related to Petunia. In wild type Brunfelsia flowers fade after bud opening ^7^, similar to *ph3*, or *ph4* mutants in Petunia (Supplementary Fig. 8a). To assess whether fading in *B. calycina* might involve FA, we identified the *BcalFA* homologue (Fig. 7a; Supplementary Fig 2). In Brunfelsia, *BcalFA* is highly expressed in petals, but hardly in leaves (Supplementary Fig. 8b), while expression of a *35S:BcalFA* transgene in *P. hybrida* V74 (fa) restored the fading phenotype, with similar efficiency as PhybFA (Supplementary Fig. 8c). Together this suggests that FA is likely to be involved in fading in (wild type) Brunfelsia flowers

**Figure 7.**
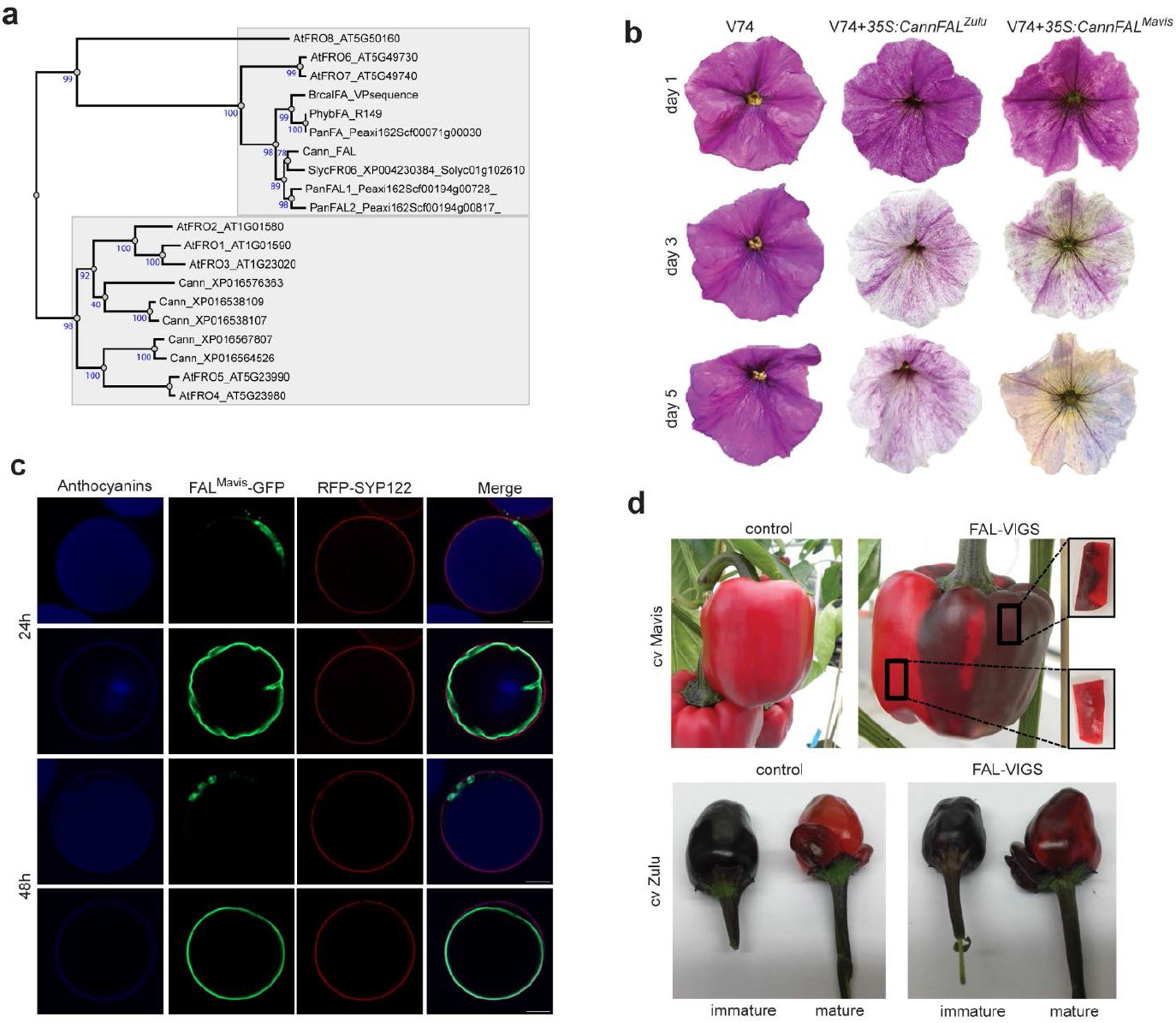
Silencing of FA in pepper results in purple mature fruits. **(a)**. Phylogenetic tree of proteins of the FRO family in *Capsicum annuum* (Ca). This tree is the result of 300 bootstraps. Numbers indicate percentage of bootstraps. **(b)**. The fading phenotype is restored in the petunia *fa* mutant line V74 by the expression of the FA homologue from pepper (*CannFAL*) from a *35S:CaFA* construct. Both the allele of the Zulu and the Mavis cultivar induce loss of anthocyanin pigmentation in petals. **(c)** Subcellular localization of the FAL^Mavis^-GFP fusion protein in wild type petal protoplasts, one or two days after transformation. RFP-AtSYP122 (visible in red) is used as marker of the plasma membrane, anthocyanins the lumen of the CV of epidermal cells are visible in blue. The size bars equal 10 μm. **(d)** Phenotype of fruits of the cultivars Mavis and Zulu and the effect of VIGS for the endogenous *FAL* gene on the permanence of anthocyanins in the peel and flesh.

In many fruits anthocyanins fade away during the maturation, causing loss of nutritional value^34^ . In some pepper cultivars, fruits have a strong purple color due to anthocyanin accumulation early in development, which is lost during maturation for which fully ripe fruit is white or only colored by carotenoids^35,36^.The *Capsicum annuum* genome appears to lack a cognate FA ortholog, as the Capsicum protein with most similarity to FA (XP_016538897) groups with FAL1 and FAL2 (Fig. 7a) and because encoding genes resides in a region that is (micro) syntenic to the Petunia *FAL1* region. Hence, we designated this gene *CannFAL1. CannFAL* homologs from two varieties of pepper (Mavis and Zulu) with strongly fading fruits are both able to complement the *fa* phenotype in petunia V74 (Fig. 7b) and GFP fusions these proteins show a similar intracellular localization as FA-GFP when expressed in *Petunia* petal protoplasts (Fig. 7c and Supplementary Fig. 9). Silencing via VIGS (Virus Induced Gene Silencing) of the *CannFAL* gene in Mavis and Zulu plants resulted in retention of anthocyanins in mature fruits (Fig. 7d and Supplementary Fig. 10a-b). These results indicate that the loss anthocyanins and the purple color during maturation of pepper fruits involves a similar mechanism as in fading Petunia flowers. Moreover, they suggest that varieties with purple mature fruits may be obtained by classical breeding for loss of function mutations, instead of transgenic approaches

## Discussion

To understand how the MBWW complex AN1-AN11-PH4-PH3 promotes the stability of anthocyanins in the central vacuole, we identified FADING showing that it encodes a vacuolar membrane ferric reductase oxidase (FRO) protein that promotes anthocyanin degradation and thereby “fading” of the petal color.

FROs play diverse roles in the plant cells, as mirrored by their expression in distinct tissues and cell types, and their localization in different cell compartments^37,38^. AtFRO2 is a plasma membrane-based ferric reductase and the paralogs AtFRO4/AtFRO5 cupric reductases thought to be in the plasma membrane that reduce iron and copper for uptake (as Fe^2+^ and Cu^+^) from the soil ^21–23^. FRO3 and FRO8 are expressed in leaf veins and predicted to be mitochondrial proteins^23^. The FA homolog AtFRO7 was localized in chloroplasts^32^ in agreement with the reduced Fe content of the chloroplasts in *fro7* mutants. Our data confirm that AtFRO7 is indeed targeted to chloroplast, and show that in petals cells, which lack chloroplasts can also be targeted to vacuolinos, just like FA. AtFRO6 is not targeted to the plasma membrane^32^, but to vacuoles/vacuolinos (Supplementary Fig. 6) just as the colocalizing AtAHA10 protein^39^.

*FA* is expressed in a broad range of tissues where it is targeted to the tonoplast of the CV to trigger the degradation of any anthocyanins in there. However, in epidermal petal cells FA is upheld in vacuolinos and anthocyanin in the central vacuole are protected from degradation. While vacuolinos are an attractive model to identify proteins involved in membrane trafficking^9^, their biological function remained unclear. Our results suggest that vacuolinos act as gatekeepers in vacuolar trafficking that allow some proteins to move on to the CV, while retaining others.

The arrest of FA-GFP, FAL1-GFP, AtFRO6-GFP or AtFRO7-GFP, is associated with release of GFP fluorescence from the membrane into the vacuolino lumen. For several reasons we consider it unlikely that this is due to release of the GFP tag by proteolysis. First, immunoblot analysis did reveal not massive cleavage products. Second, expression of N-terminally tagged GFP-FA also results in GFP fluorescence appearing the vacuolino lumen. Third, analysis of the membrane topology of FRO2 indicates that the N- and C-termini of the protein are on the cytoplasmic side of the membrane. Together, this suggests that the observed internalization of the GFP fluorescence must involve a more complex endocytic process than proteolytic cleavage.

The mechanism by which FA triggers anthocyanin degradation is subject of current research. The dark side of FRO activity is the production of ROS (Reactive Oxygen Species) because Fe participates in electron transfer reactions that generate radicals^40^. One possibility is FA activity generates in the CV lumen Fe^2+^ ions that by returning to Fe^3+^ generate ROS that are scavenged by anthocyanins (antioxidants), resulting in degradation of the pigment molecules. Another possibility is that the produced Fe^2+^ is needed for activity of a Fe-binding enzyme involved in anthocyanin degradation. In vitro work showed a class III peroxidase implicated in the degradation of anthocyanin in developing *Brunfelsia* petals. Whether this peroxidase has a role in *in vivo* fading and operates in concert with BrcalFA – which triggers anthocyanin degradation in petunia petals – remains to be established.

Increased accumulation of anthocyanin pigments in specific plants tissues is a desired trait in breeding of ornamentals and food crops for the (pretty) novel color and the associated health promoting effects^6,41^. Crops with increased anthocyanin accumulation in edible parts can be easily generated though the ectopic expression of transcription factors^6,42^, but are not suitable for marketing because they are transgenic. Our results suggest that by focusing on the FA anthocyanin degradation pathway may an alternative based on loss of function mutations that can be isolated by conventional strategies.

## Online methods

### Plant material

All petunia lines originate from the petunia collection of the University of Amsterdam. Family numbers (e.g., R2057) are referred to all individuals originating from one single pollination, then individuals get a unique number (e.g., R2057-1).

M1×V30 is an F1 hybrid between the peonidin producing line M1 (*FA PH4 PH3*) and the malvidin line V30 (*fa PH4 PH3*). R159 is an unstable *ph4* mutant line isogenic with the cyanidin wild type line R27. For analysis of the effect of the *ph3* mutation on the expression of the *FA* gene and the fading phenotype we used the segregating progeny of the selfing of a R143 heterozygous individual (*PH3 ph3*). The *ph1* (M1020, *ph1*^*V23/V23*^) mutant was generated by crossing (V23×V30) × V23. In this population we identified isogenic mutant and wild type plants. The *Brunfelsia calycina*, plants have been obtained from seeds kindly provided by the Solanaceae collection and Gene Bank of the University of Nijmegen (which was recently discontinued).

### Mass Spectrometry and data analysis

Samples (small pools of corollas from specific genetic background and specific developmental stages) were nitrogen frozen, powdered with mortar and pestle, stored at -80°C and, just before the analysis, extracted for 30 minutes with 6 volumes (w/v) (for HPLC-ESI-IT and HPLC-DAD) or 10 volumes (w/v) of methanol/HCl 99/1 (for UPLC-ESI-QTOF), diluted with 8 volumes (v/v) (for HPLC-ESI-IT) or 100 volumes of water (V/V) (for UPLC-ESI-QTOF), filtered and analysed. Sample analysis and data processing were performed as described before^43,44^. For metabolite identification “in house” libraries of metabolite structures and of spectra of authentic standards were used. Multivariate statistical analysis was performed with Simca v. 13.0.3.0 (Umetrix AB, www.umetrix.com). A permutation test with 200 permutations and the ANOVA test were used to validate the OPLS-DA (Orthogonal Projection to Latent Structure-Discriminant Analysis) model. Modelling of AC fragmentation was performed with Chem Sketch v.2018.2.1 (ACD Labs, www.acdlabs.com).

### RNA extraction and RT-qPCR

Total RNA was isolated from petunia tissues using the TRIzol reagent (Thermo Fisher Scientific). Real-time RT-PCR was performed with an iTaq Universal Syber Green kit (Bio-Rad) using primers listed in Supplementary Table 2 and an Applied Biosystems™ QuantStudio 3 Real-Time PCR Systems and QuantStudio Design Analysis Software.

### RNAseq data

SRA data^28,29^ of of *P. axillaris, P. inflata*, a F1 progeny and *P*. exserta, were trimmed (trimmomatic) and mapped (hisat2) to their reference genome. The alignment files (bam-files) were used to get a de novo annotation (stringtie) and expression values (featureCounts and DESeq2). All public RNA-seq data are deposited in the NCBI Sequence Read Archive under BioProjects SRA: PRJNA344710, PRJNA674380, PRJNA482765.

### Amplification of *PhFA* and its homologs from different species

With combinations of primers based on the petunia sequence and sequences retrieved from a public transcriptome (Bioproject accession number PRJNA229456) of *Brunfelsia calycina* (which flowers show fading)^31^ we could amplify the full sequence of the homolog from this species. For Arabidopsis FRO6 and FRO7 we designed primers based on published sequences (AT5G49730.1 and AT5G49740.1, respectively) in The Arabidopsis Information Resource (TAIR).

### Transposon flank sequencing

Transposon flanking sequences were amplified using a modification of transposon-display as described before^45^. Genomic DNAs isolated from petals of putative non-fading mutants and nine fading siblings that originated from the same pollination/seed capsule were digested with *MunI* and *MseI/BfaI*, ligated to a biotinylated *MunI* adapter and an *MseI/BfaI* adapter. Ligation products which were subsequently enriched by magnetic particles. Further enrichment of the target fragments by preamplification were conducted with primer pairs presenting in the adapters. Then reamplification of the candidate transposon-flanking sequences by PCR with the *dTPH1* invert repeats primer bearing different barcodes for different samples. Finally, pooling all the barcoded-flanks and introducing the illumine adapter to barcoded-flanks, which were sent out for Illumina sequencing.

### Constructs

Coding sequence (CDS) or genomic DNA (gDNA) sequences of *FA, FAL, FRO6, FRO7*, were amplified from (wild-type) cDNAs or gDNAs with specific primers (as reported in Supplementary Table 2) containing attB1 and attB2 sites using Phusion High-Fidelity DNA Polymerase. Purified PCR product was subsequently cloned into Gateway pDONR221 donor vector as an entry clone by GatewayBP recombination reaction (Invitrogen Clonase Gateway BP Clonase II EnzymeMix). Finally Gateway LR recombination reaction (Invitrogen Clonase Gateway LR Clonase II Enzyme Mix) was performed with entry clone and multiple destination vectors^10^ to make functional plasmids. *35S:PH5-GFP* and *35S:RFP-SYP122* were previously generated and described in previous works^17,33^.

The VIGS construct for the silencing in pepper of the endogenous *CannFA* gene was obtained using a target region from nucleotide 113 to 384 of the FA gene (coding sequence). This fragment was cloned in a vector derived from TRV2 plamid^46^. Twelve plants were obtained and silencing was observed in all plants.

### Anthocyanin content determination in pepper fruits

Fruits were weighed, ground in liquid N_**2**_, and after, addition of five volumes (based on weight) of extraction buffer (45% methanol, 5% acetic acid) centrifuged at 12,000 x g for 5 min at room temperature. The supernatant was then to a new tube and again centrifuged at 12,000 x g for 5 min. The absorption at 530nm was then determined with a spectrophotometer.

### Transient expression in protoplasts and confocal microscopy

Protoplasts isolated from the limb of opened flowers of different were transformed with plasmids of interest as described before^47,48^. Transfected cells were observed and imaged under Zeiss LSM510 confocal microscope using a 40×/1.2 water objective at 24 or 48 hours after transformation. GFP was excited using a 488 nm laser paired with a 530/30 nm bandpass filter. RFP was excited using a 561 nm laser paired with a 582/15 nm bandpass filter.

### Phylogenetic analysis

Phylogenetic trees were inferred by PhyML using sequences that had been aligned with MUSCLE and cured with BGME using online tools at https://ngphylogeny.fr.

### Measure of *Fe* redox activity in plant petals

Fe reducing activity of petal extracts was determined as described previously^49^, with some modifications. Opened flowers were collected for each plant genotype and ground in liquid nitrogen. Anthocyanins were removed by the addition of a teaspoon of Dowex (Dowex 1X2 chloride form, 50-100 mesh, Sigma-Aldrich) during grinding. The resulting powder was suspended in ice-cold buffer (50 μM MES at pH 5.5 and 40 nM CaSO_4_), centrifuged for 3 minutes at 3214 g at 4 °C. The supernatants were transferred to new tubes, together with Fe(III)-EDTA, BPDS (Bathophenanthrolinedisulfonate, Sigma-Aldrich) and NADH (final concentrations of 10 μM, 1 mM, and 4 μM, respectively). Samples were then kept in the dark and production of Fe(II)-BPDS was calculated from the extinction at 535 nm, using an extinction coefficient of 22.14 mM^-1^cm^-1^ .

## Supporting information

DataSheet 1

## Acknowledgements

The authors are thankful to Pieter Hoogeveen and Floris Marsman for the care of the petunia plants and to Bets Verbree for the administration of the petunia collection. The authors also thank Stefania Ceoldo for her technical support at the Department of Biotechnology, University of Verona. We would have never been able to achieve these results without the help and assistance of the group of Prof. D. Gadella of the Amsterdam Centre for Advance microscopy of the UvA (https://sils.uva.nl/content/research-groups/molecular-cytology/lcam/van-leeuwenhoek-centre-for-advanced-microscopy.html).

