## Supplementary info for "Role of the metallo-reductase FADING and vacuolinos in anthocyanin degradation in flowers and fruits"

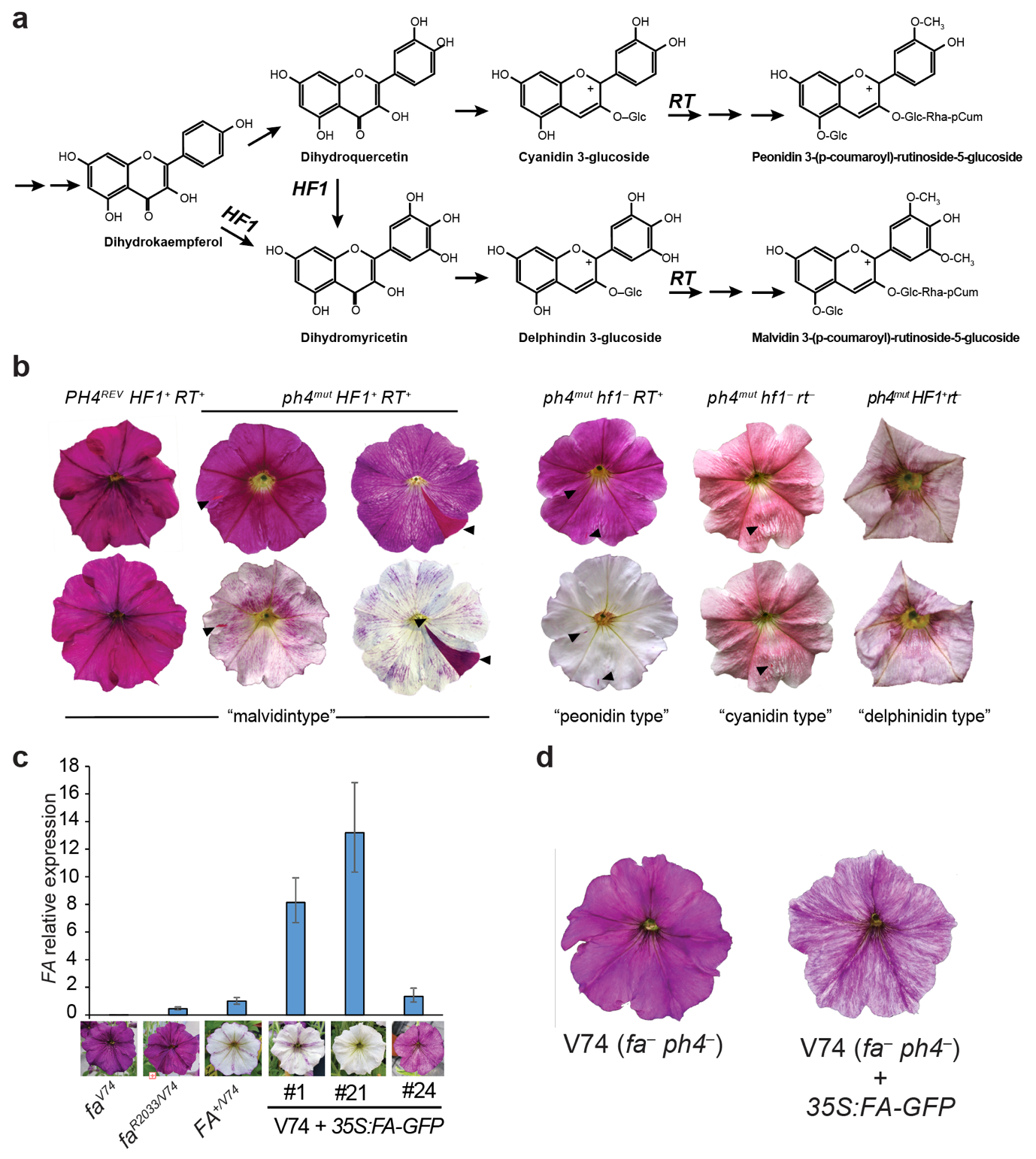

**Supplementary Figure 1. Genes affecting flower color fading.**

**(a)** Simplified representation of anthocyanin synthesis showing the role of the genes HF1, encoding flavonoid 3’5’ hydroxylase and RT encoding anthocyanin rhamosyltransferase.

**(b)** Flowers from F2 progeny of lines V74 (*fa, ph4, Hf1, RT*) and R159 (F*A, ph4^mut^, hf1 rt*), shortly after bud opening (top row) and 6 days later (bottom row). The flower shown contain ra dominant FA allele, and segregate for *HF1*, *RT* and revertant *PH4^REV^* allele The arrowheads indicate *PH4* reversion spots in which fading does not occur. Notice that flowers accumulating delphinidin do not unfold their petals completely. **(c)** Phenotype of the flowers of individual *35S:FA* (V74 + *35S:FA*) compared to the untransformed controls (V74) and the R2033 plants carrying the footprint allele of *fa* (*fa^R2033-1^*) or the wild type *FA* allele (*FA^R149^*).The expression level of the *FA* transcript correlates with the degree of pigmentation loss. **(d)** The GFP fusion of the FA protein (FA-GFP) is also able to restore the fading phenotype in the *fa* mutant V74.

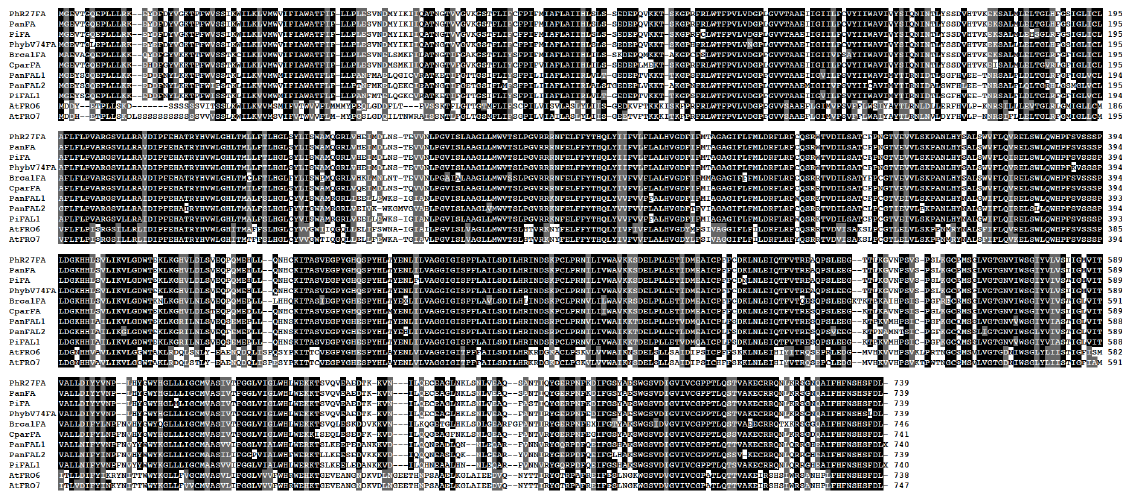

**Supplementary Figure 2.** Alignment of FA and FAL proteins from *Petunia hybrida* R27 (PhR27), *Petunia axillaris* N (Pan), *Petunia inflata* (Pi), *Petunia hybrida* V74 (PhybV74),and FA homologs from *Brunfelsia calicina* (Brcal), *Calibrachoa parviflora* (Cpar), and *Arabidopsis thaliana* (At).

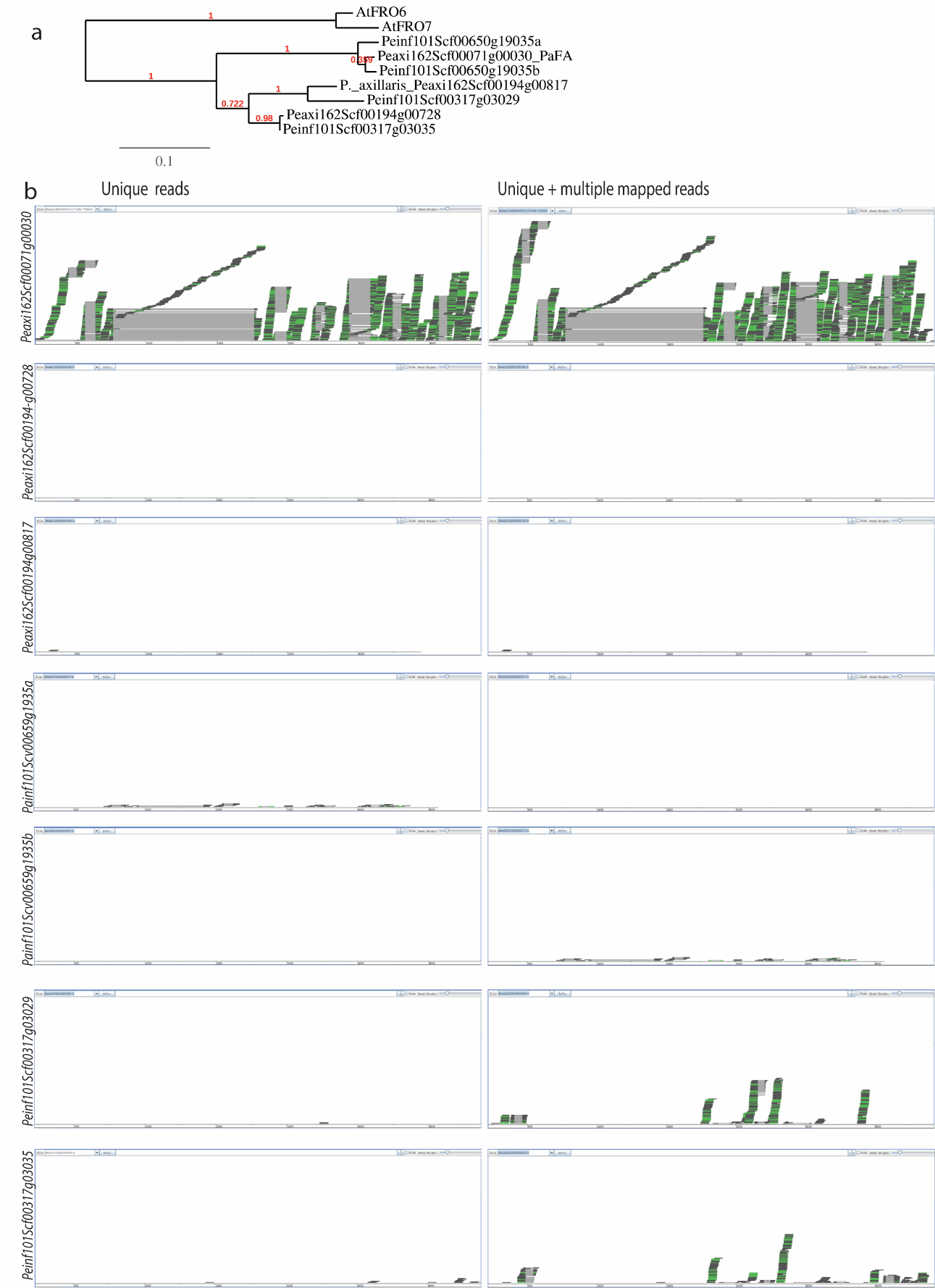

**Supplementary Figure 3.** **Expression of *FA* and *fa* alleles. (a)** *FA* and *FAL* genes in *P. axillaris* and *P. inflata*. **(b)** Mapping to *FA* and *FA-like* genes of the reads generated from RNA-seq of petals of F1 (*P.axillaris* × *P.inflata*) plants. Unique reads for each gene are shown in the first column, and reads that map to more than one gene have been discarded; in the second column, in addition to unique reads, multiple mapped reads are kept and reported in the graph.

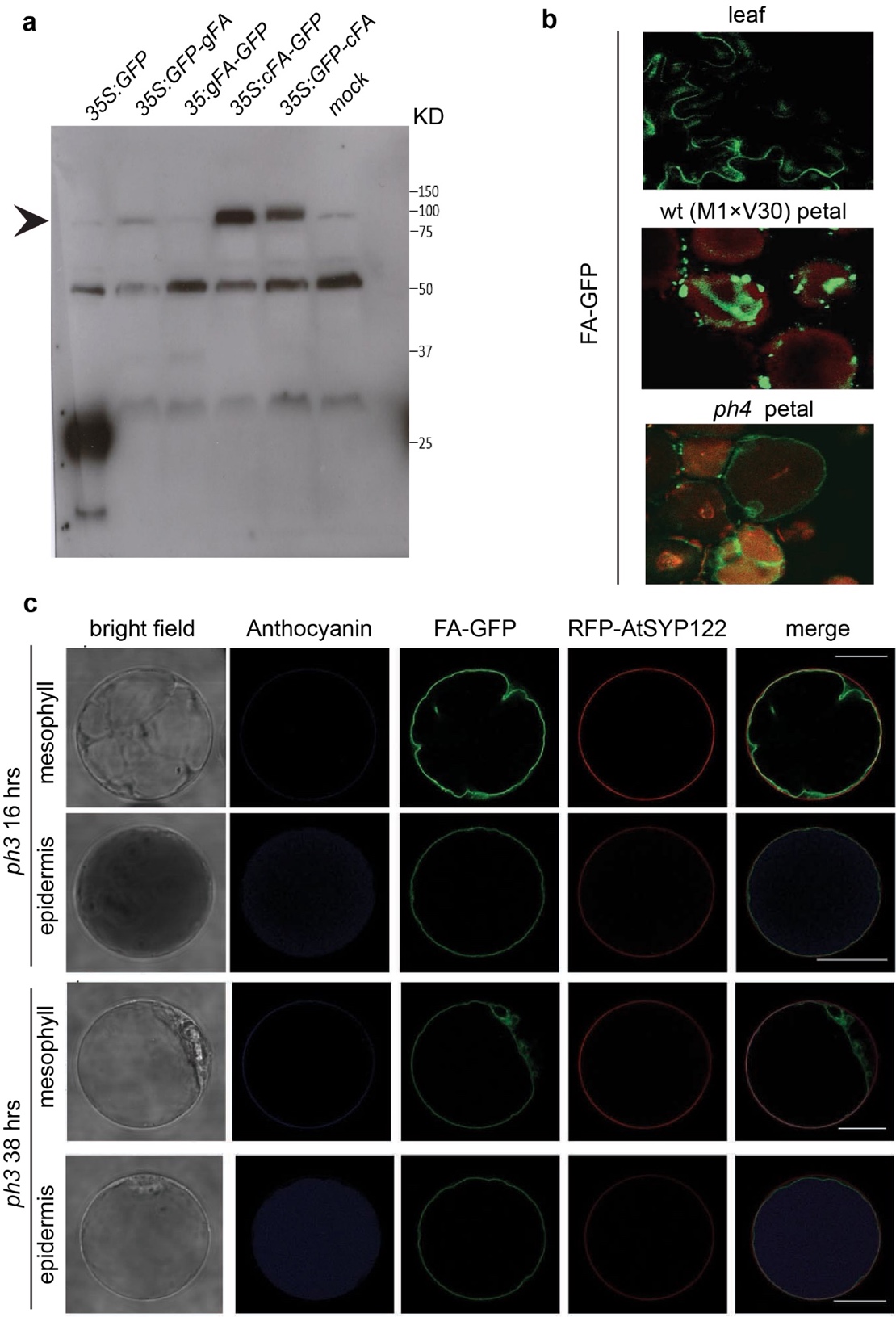

**Supplementary Figure 4.** **Subcellular localization of FA protein**. **(a)** Western blot analysis of proteins extracted from wild type petunia petals infiltrated with Agrobacterium harboring the constructs *35S:GFP,* *35S:FA-GFP*, *35S:GFP-FA* or water (mock) . The FA protein was expressed from the cDNA sequence (cFA) or the genomic fragment containing the sequence from ATG to STOP codon (gFA). Total proteins were extracted from petals 24 hours after infiltration. The arrow points out the expected bands for fusion protein. **(b)** Leaves and petals infiltrated with Agrobacterium harboring the *35S:cFA-GFP* construct. **(c)** Localization of FA-GFP (in green) in petal protoplasts from *ph3* mutant petals. RFP-SYP122 (visible in red) is used as marker of the plasma membrane, anthocyanins in the lumen of the CV of epidermal cells are visible in blue. The size bars equal 20 μm.

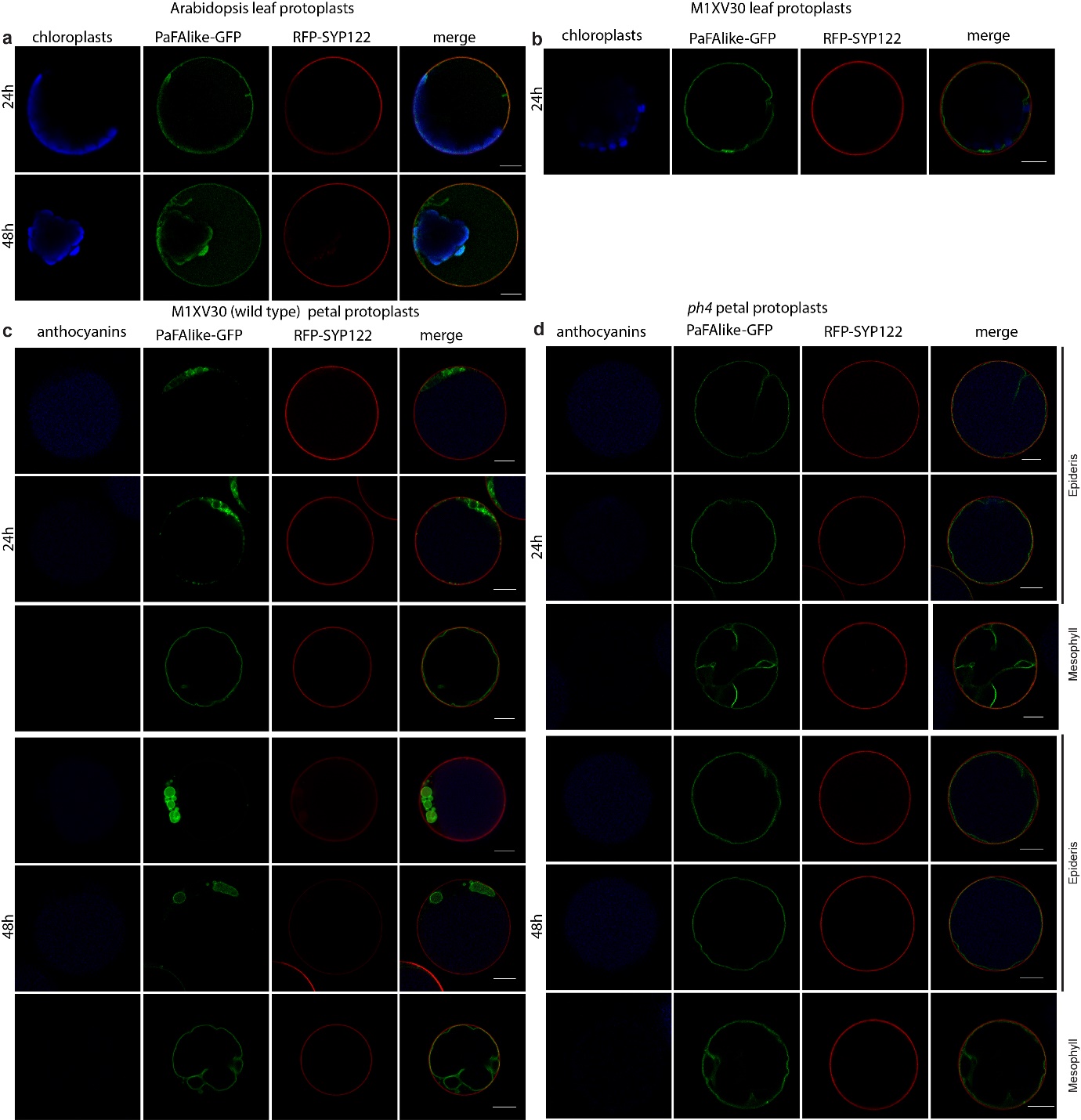

**Supplementary Figure 5**. **Subcellular localization of petunia PaFA1-like (Peaxi162Scf00194g00728) in transiently transformed protoplasts.** **(a-c)** Localization of PanFA1-like in protoplasts from wild type Arabidopsis leaves **(a)**, wild type petunia leaves **(b)**, wild type petunia petals **(c)**, *ph4* petunia petals **(d)** In protoplasts from petals, one or two days after transformation. RFP-AtSYP122 (visible in red) is used as marker of the plasma membrane, anthocyanins the lumen of the CV of epidermal cells are visible in blue. The size bars equal 10 µm.

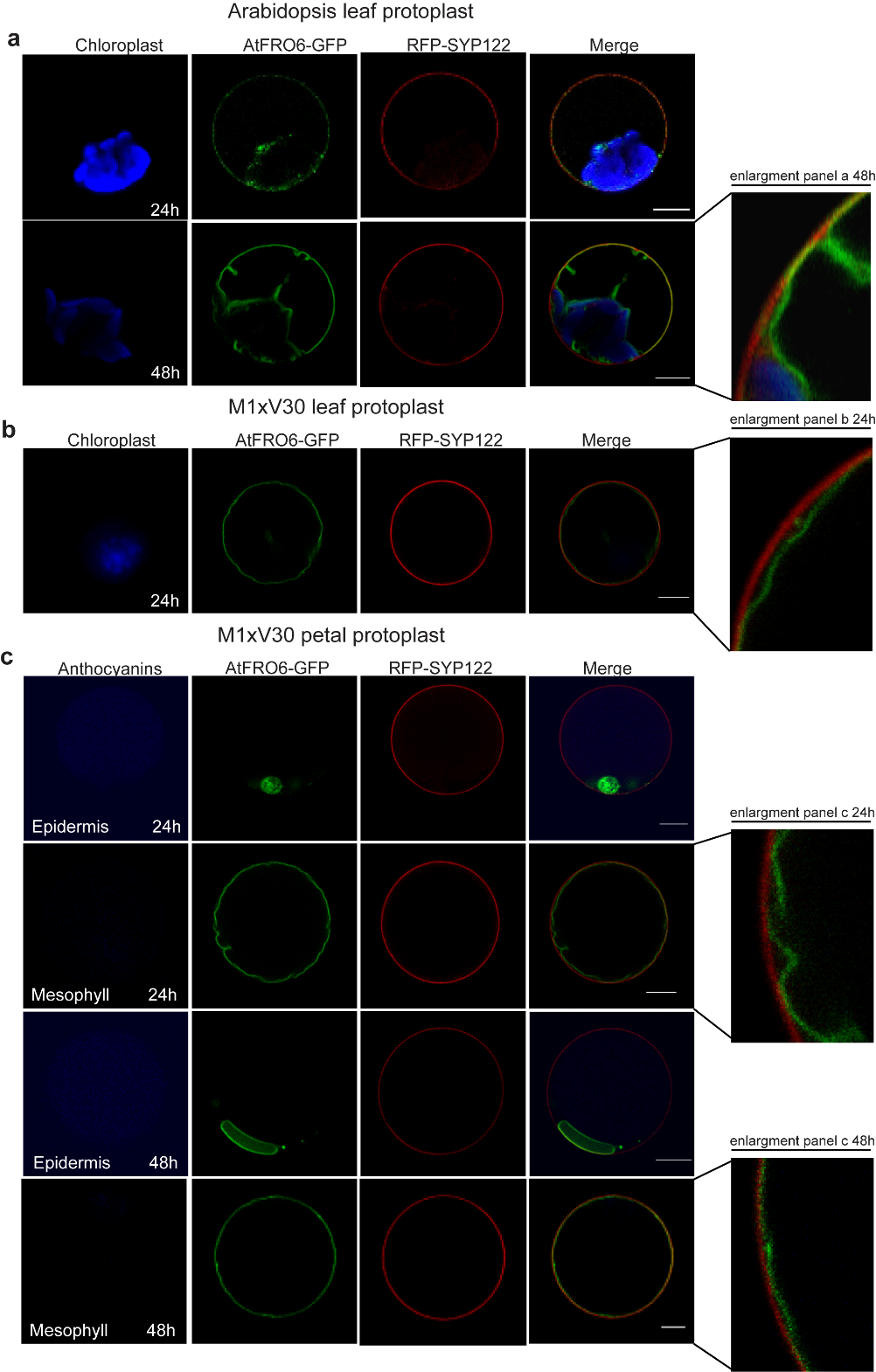

**Supplementary Figure 6. Subcellular localization of Arabidopsis FRO6 in transiently transformed protoplasts. (a-c)** Localization of AtFRO6-GFP in protoplasts from wild type Arabidopsis leaves **(a)**, wild type petunia leaves **(b)**, wild type petunia petals **(c)** one or two days after transformation. RFP-AtSYP122 (visible in red) is used as marker of the plasma membrane, anthocyanins in the lumen of the CV of epidermal cells are visible in blue. The size bars equal 10 µm.

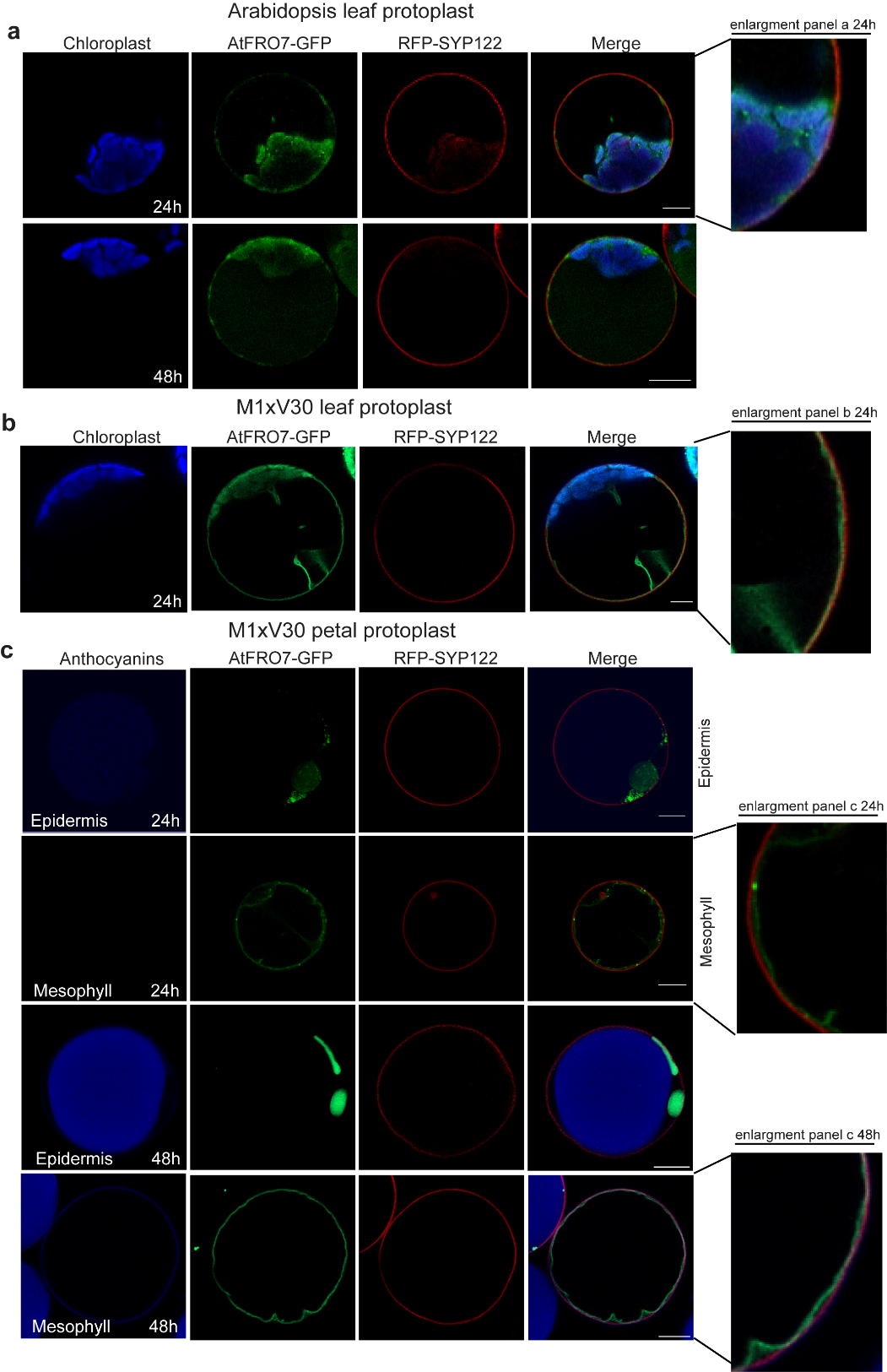

**Supplementary Figure 7. Subcellular localization of Arabidopsis FRO7 in transiently transformed protoplasts. (a-c)** Localization of AtFRO6-GFP in protoplasts from wild type Arabidopsis leaves (a), wild type petunia leaves (b), wild type petunia petals (c) one or two days after transformation. RFP-AtSYP122 (visible in red) is used as marker of the plasma membrane, anthocyanins the lumen of the CV of epidermal cells are visible in blue. The size bars equal 10 µm.

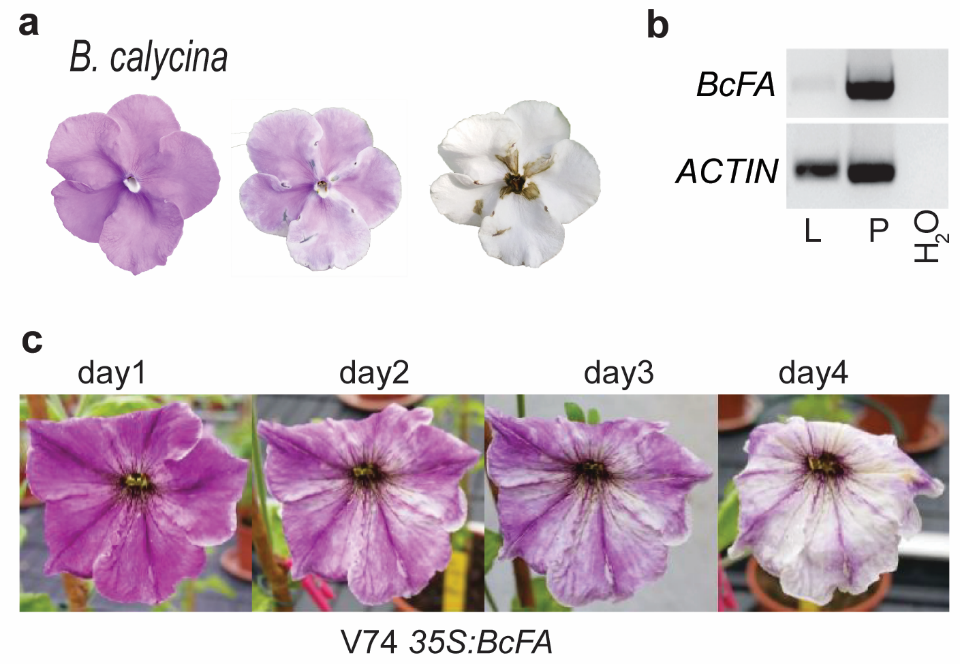

**Supplementary Figure 8.** **Complementation of the fa phenotype with the FA gene from Brunfelsia calycina.** **(a)** The same flower was photographed for 4 days starting from bud opening (day 1). **(b)** RT-PCR of the transcripts for the *BcFA* gene in leaves and petals of *Brunfelsia*. **(c)** Flowers from a transgenic petunia V74 (*ph4 fa*) plant in expression of a 35SBCFA transgene restores fading.

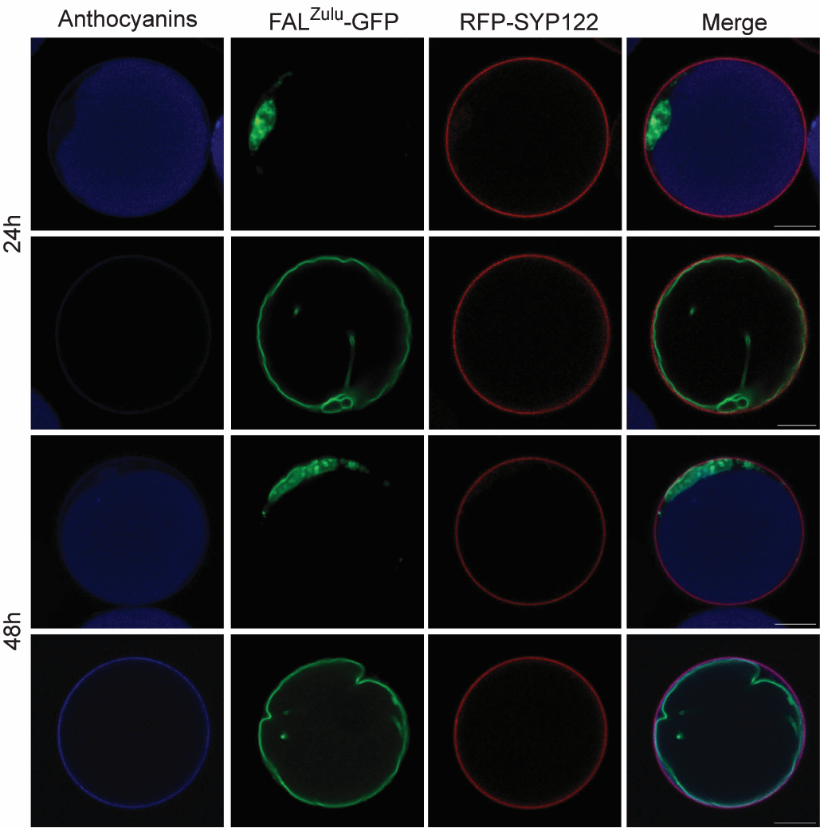

**Supplementary Figure 9.** Subcellular localization of FAL^Zulu^-GFP fusion protein in transiently transformed wild type petal protoplasts, one or two days after transformation. RFP-AtSYP122 (visible in red) is used as marker of the plasma membrane, anthocyanins the lumen of the CV of epidermal cells are visible in blue. The size bars equal 10 µm.

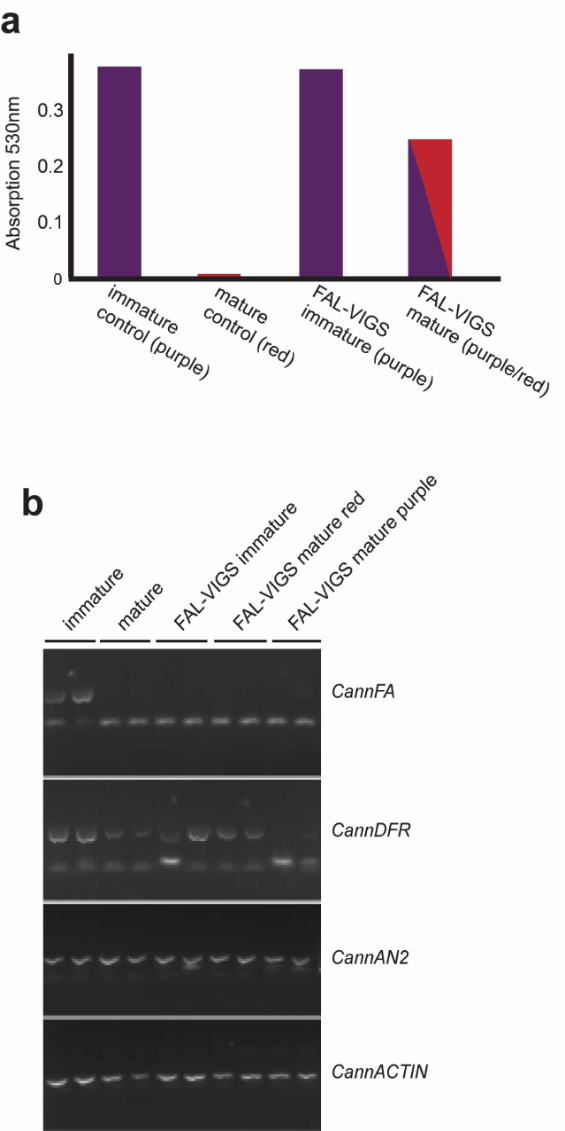

**Supplementary Figure 10**. **Silencing of FA in pepper results in purple mature fruits.** **(a)** Concentration of anthocyanins in mature and immature pepper fruits (variety Zulu). When the FA gene was silenced, (FA expression reduced), high anthocyanins concentration in both the immature and mature fruits is maintained. **(b)** Expression (RT-PCR) of the *CannFAL* gene in control and FAL-VIGS fruits of Zulu cultivar, compared to the expression of the structural anthocyanin gene *CannDFR*, the anthocyanin regulator *CannAN2* and the housekeeping gene *CannActin*.

| **Supplementary Table 1. Putative anthocyanin fragments detected in petals after fading.** | | | | | | | | |
| --- | --- | --- | --- | --- | --- | --- | --- | --- |
| **HPLC-ESI-IT^1^** | | | | **UPLC-ESI-QTOF^2^** | | | | **putative identification** |
| **rt** | **m/z(neg)** | **ms/ms(neg)** | **ms3(neg)** | **m/z(-) expected** | **m/z(-) detected** | **elemental composition** | **mass error** |  |
| 4.36314 | 330.687 | 169, 161, 154, 139, 125, 113 |  | 331.0665 | 331.0661 | C_13_H_16_O_10_ | 1.208 | trihydroxybenzoic acid hexose RING B of delphinidin isomer 1 |
| 5.70969 | 330.694 | 169, 167 | 167(153,154, 139) | 331.0665 | 331.0674 | C_13_H_16_O_10_ | -2.718 | trihydroxybenzoic acid hexose RING B of delphinidin isomer 2 |
| 19.5551 | 470.701 | 309, 187, 157 | 309(119, 144, 163,187) | 471.1502 | 471.1505 | C_21_H_28_O_12_ | -0.637 | 1 coumaroyl rutinose isomer 1 |
| 20.5373 | 470.715 | 411, 381, 351, 309 | 309(203, 187, 163,145) | 471.1502 | 471.1512 | C_21_H_28_O_12_ | -2.122 | 1 coumaroyl rutinose isomer 2 |
| 24.3736 | 808.712 | 647, 501, 453, 351 | 647( 501,269); | 809.2140 | 809.2123 | C_36_H_42_O_21_ | 2.101 | anthocyanin (coumaric acid) 3 O rutinose 5 O glucose – (B RING + H_2_O) isomer 1 |
| 26.0199 | 808.728 | 647, 501, 453, 351 | 647(501,453,351, 339) | 809.2140 | 809.2125 | C_36_H_42_O_21_ | 1.854 | anthocyanin (coumaric acid) 3 O rutinose 5 O glucose – (B RING + H_2_O) isomer 2 |
| 21.5373 | 824.733 | 663, 501, 465, 366 | 501(473,265, 235, 218, 207,193,179,166) | 825.2089 | 825.2081 | C_36_H_42_O_2_2 | 0.969 | anthocyanin (caffeic acid) 3 O rutinose 5 O glucose – (B RING + H_2_O) |
| 25.1467 | 838.758 | 355, 501, 677 | 677 (501) | 839.2245 | 839.2242 | C_37_H_44_O_22_ | 0.357 | anthocyanin (ferulic acid) 3 O rutinose 5 O glucose – (B RING + H_2_O) |

^1^For HPLC-ESI-IT the m/z feature values (negative ionization mode) and the fragmentation trees (ms/ms and MS^3^) are given.

^2^For UPLC-ESI-QTOF the high resolution detected m/z (negative ionization mode) are compared with the expected, and the mass error, in ppm, is given.

**Supplementary Table 2.** Primers used in this study.

| Purpose | Gene | Sequence 5'-3' | Ori^1^ |
| --- | --- | --- | --- |
| ***Genotyping*** | | | |
| *Fading ^R2057^* | *PhFA* | TAGGAGACTGGACCGAAAAGC | F |
|  |  | CCAACCAAACCAGACATTGGAC | R |
| ***RT-PCR*** | | | |
|  | *PhFA* | ATGGGTGAAGTCACAGGCCA | F |
|  |  | CTTTTCTCTTTCACAGTATGTACATC | R |
|  | *BcFA* | TGAGACTGGTTTACATAAGCTTTCC | F |
|  |  | AAGGCTTGCTGTCATTAATCAGG | R |
| Reference gene | *PhACTIN* | AGATCTGGCATCATACCTTCTACA | F |
|  |  | CCMGCAGCTTCCATRCCAATCA | R |
| ***qRT-PCR*** | | | |
|  | *PhFA* | TTGATGCTGGAGCTCACAGG | F |
|  |  | GCATTGTTAGATGTCCCAGCCA | R |
|  | *PhFA* | GGCTGGGACATCTAACAATGCTTC | F |
|  |  | GTCACCCACATCAATAAACCAGC | R |
|  | *Cann_FAL* | TTGTTTCTGCCTGTTGCACG | F |
|  |  | AAGGGAAGCATGTGGCTGAA | R |
|  | *CannAN2* | AGCTTCTAGGCAACAGATGGT | F |
|  |  | TGTGGTGATCTTGAGGGCAG | R |
|  | *CannDFR* | AGCAGACTTGACCGTGGAAG | F |
|  |  | CTTCGTTCTCAGGGTCCTTGG | R |
| Reference gene | *PhRAN* | CAGGCAGGTTAAGGCAAAGC | F |
|  |  | CTAGCAAGGTACAGAAACGGC | R |
| Reference gene | *CannACTIN* | ATCCCTCCACCTCTTCACTCTC | F |
|  |  | GCCTTAACCATTCCTGTTCCATTATC | R |
| CDS | *PhFA* | ggggacaagtttgtacaaaaaagcaggctatatgggtgaagtcacaggcca | F |
|  |  | GGGGACCACTTTGTACAAGAAAGCTGGGTCTAGGTCAAAACTATGGCTGTTGAA | R (+stop) |
|  |  | GGGGACCACTTTGTACAAGAAAGCTGGGTCTAG GTC AAA ACT ATG GCT GTT GAA | R (-stop) |
| CDS | *BcFA* | ggggacaagtttgtacaaaaaagcaggctCAATGGCTGAAGTTGCAGGCCAAG | F |
|  |  | GGGGACCACTTTGTACAAGAAAGCTGGGT CTA TAG GTC AAA ACT ATG GCT GTT G | R (+stop) |
|  |  | GGGGACCACTTTGTACAAGAAAGCTGGGT TTAG GTC AAA ACT ATG GCT GTT G | R (-stop) |
| CDS | *AtFRO6* | GGGGACAAGTTTGTACAAAAAAGCAGGCTCCATGGATGATTATGAAACCCCTCTTTTG | F |
|  |  | GGGGACCACTTTGTACAAGAAAGCTGGGTAGAGATCGAAACTGTGGCTGTTG | R (-stop) |
| CDS | *AtFRO7* | GGGGACAAGTTTGTACAAAAAAGCAGGCTCCATGGATGATCATGAAACCCCTC | F |
|  |  | GGGGACCACTTTGTACAAGAAAGCTGGGTAGAGATCGAAACTGTGGCTGTTG | R (-stop) |
| CDS | *PaFA-like1* | GGGGACAAGTTTGTACAAAAAAGCAGGCTTTATGGGTGAATATTCAGGCCAAGAAC | F |
|  |  | GGGGACCACTTTGTACAAGAAAGCTGGGTAGAGGTCAAAACTGTGGCTGTTAAAATG | R (-stop) |
| gDNA | *CannFAL-Mavis* | GGGGACAAGTTTGTACAAAAAAGCAGGCTATATGGGTGAACTTTCAAGCCAAG | F |
|  |  | GGGGACCACTTTGTACAAGAAAGCTGGGTCTAGAGGTCAAAACTGTGGCTG | R (+stop) |
|  |  | GGGGACCACTTTGTACAAGAAAGCTGGGTGAGGTCAAAACTGTGGCTGTTG | R (-stop) |
| gDNA | *CannFAL-Zulu* | GGGGACAAGTTTGTACAAAAAAGCAGGCTATATGGGTGAACTTTCAAGCCAAG | F |
|  |  | GGGGACCACTTTGTACAAGAAAGCTGGGTCTAGAGGTCAAAACTGTGGCTG | R (+stop) |
|  |  | GGGGACCACTTTGTACAAGAAAGCTGGGTGAGGTCAAAACTGTGGCTGTTG | R (-stop) |
| RNAi | *PaFA* | GGGGACAAGTTTGTACAAAAAAGCAGGCTGCAAACCCGAGTAATTGTTTCATAG | F |
|  |  | GGGGACCACTTTGTACAAGAAAGCTGGGTTTTTGAAACTCAGGAAATTCCATTAACAG | R |

^1^ F indicates a forward orientation of the primer, relative to the orientation of the gene, and R reverse orientation.

Sequence marked with green and red represent *att*B1 and *att*B2 respectively.

*Ph, Petunia hybrid; Pa, Petunia axillaris; Bc, Brunfelsia calycina; At, Arabidopsis thaliana; Ca, Capsicum annuum*
